# An adhesion code ensures robust pattern formation during tissue morphogenesis

**DOI:** 10.1101/803635

**Authors:** Tony Y.-C. Tsai, Mateusz Sikora, Peng Xia, Tugba Colak-Champollion, Holger Knaut, Carl-Philipp Heisenberg, Sean G. Megason

**Author notes:** Max Planck Institute of Biophysics Frankfurt am Main, Germany (Mateusz Sikora).

## Abstract

An outstanding question in embryo development is how spatial patterns are formed robustly. In the zebrafish spinal cord, neural progenitors form stereotypic stripe-like patterns despite noisy morphogen signaling and large-scale cellular rearrangement required for tissue growth and morphogenesis. We set out to understand the mechanisms underlying this patterning robustness. Our adhesion assays revealed a preference for three neural progenitor types to stabilize contacts with cells of the same type. Genetic analysis uncovered a three-molecule adhesion code, composed of N-cadherin, Cadherin 11, and Protocadherin 19, with unique gene expression profiles for each cell type. Perturbation of the adhesion code results in loss of homotypic preference *ex vivo* and patterning errors *in vivo*. Both the cell fate and adhesion code are co-regulated by the common upstream morphogen signal Shh. We propose that robust patterning in tissues undergoing morphogenesis results from a previously unappreciated interplay between morphogen gradient-based patterning and adhesion-based self-organization.

## INTRODUCTION

An outstanding question in embryo development is how spatial patterns of distinct cell types arise reproducibly. The classic French Flag model posits that morphogen gradients form across a naïve and static field of cells to provide positional information to specify patterned cell fates (Wolpert, 1969). However, it is unusual to see morphogen gradients across static fields of cells during embryo development. Tissue development generally involves significant cell proliferation and migration to change the overall tissue size and shape, a set of processes collectively termed morphogenesis (Hogan, 1999; Kicheva et al., 2014; Thompson, 1992). Both cell division and migration can lead to the disruption of existing positional information and scrambling of cell-cell neighbor relationships (Godard and Heisenberg, 2019; Guillot and Lecuit, 2013). Although the French Flag model has many important features, it does not explain how patterns can form reliably despite concurrent morphogenetic movement.

One potential mechanism for coordinating patterning and morphogenesis is through the regulation of cell-cell adhesion. Almost 50 years ago, Steinberg proposed the “Differential Adhesion model” (Steinberg, 1970), postulating that cells of different types have different adhesive affinities, allowing them to sort into cohesive domains. Subsequent studies have confirmed that this differential cell sorting exists (Foty et al., 1996), and identified some of the complex molecular mechanisms involved. The stability of cell contacts is governed by the tension between the faces of the cell contact, which is determined by a combination of cell adhesion and cortical tension (Brodland, 2002; Maître et al., 2012). Weaker adhesion and stronger contractility at the cell contact results in higher interfacial tension and less stable contact. At the molecular level, when adhesion molecules such as cadherins bind in trans, they can lower interfacial tension to stabilize cell contact by directly increasing adhesion, or signaling to actomyosin cytoskeleton to reduce contractility (Maître and Heisenberg, 2013; Maître et al., 2012). Sorting of mixed cell types can occur when homotypic contacts exhibit lower interfacial tension compared to heterotypic contacts (Brodland, 2002). This can be achieved by different cell types expressing different quantities of a single adhesion molecule, or expressing different adhesion molecules (e.g., different types of cadherins) that are mutually incompatible. Several types of adhesion molecules (e.g., cadherins and protocadherins) have been widely observed to form spatial patterns in restricted regions within tissues, especially within the vertebrate nervous system (Kimura et al., 1996; Price et al., 2002; Redies, 1995; Suzuki et al., 1997). However, differential adhesion does not explain the generation of oriented or positioned patterns: adhesion-mediated sorting simply yields homotypic clumps of cells. It is an open question whether and how morphogen-gradient-based patterning affects adhesion-mediated cell sorting within a tissue.

The zebrafish spinal cord is an ideal model system to investigate how precise spatial patterns form during morphogenesis. At the end of gastrulation, the future spinal cord tissue starts as a sheet-like structure called the neural plate. The neural plate spans a width of 400 µm along the medial-lateral and anterior-posterior axes but is as thin as one to two cells along the dorsal-ventral axis. Driven by convergent extension movement, neural progenitor cells migrate dorso-medially to form the neural keel, resulting in thickening of the tissue along the dorsal-ventral axis at the expense of shrinkage of tissue width along the medial-lateral axis. By the time this tissue becomes the neural tube, its dimensions have changed from ∼400 µm in width and ∼20 µm in height to ∼80 µm in width and ∼100 µm in height (Schmitz et al., 1993). This tissue movement is driven by cell intercalation and polarized cell divisions, resulting in the frequent changes of neighboring cells (Clarke, 2009; Tada and Heisenberg, 2012).

Concurrent with this morphogenetic movement, neural progenitor cells adopt distinct fates and form stereotypic spatial patterns that are highly conserved across vertebrate species. At the neural tube stage, thirteen different stripe-like domains can be distinguished along the ventral-to-dorsal axis, each expressing a unique combination of transcription factors (Dessaud et al., 2008; Jessell, 2000). The neural progenitor cells in these domains will eventually differentiate into distinct types of neurons with different sensorimotor functions in the spinal cord (Goulding, 2009; Goulding and Pfaff, 2005). The tissue is instructed by opposing gradients: Sonic hedgehog (Shh) is secreted by the notochord and floor plate (Echelard et al., 1993; Krauss et al., 1993). At the neural plate stage, the floor plate is positioned medially within the neural plate while the notochord is beneath it, resulting in a medial-to-lateral gradient of Shh. At the neural tube stage, both structures are positioned at the ventral side of the spinal cord, resulting in a ventral-to-dorsal gradient of Shh (Chamberlain et al., 2008; Yamada et al., 1993). On the other side of the tissue, Bone Morphogen Protein (BMP) and Wnt are secreted from the epidermal ectoderm and roof plate cells, which are initially located at the lateral side of the neural plate and later at the dorsal side of the neural tube (Chizhikov and Millen, 2005). The specification of different neural progenitor types depends on the dosage of these secreted morphogens, making the spinal cord a textbook example of the French Flag model (Dessaud et al., 2007; Ericson et al., 1997; Gilbert, 2013).

Through *in toto* imaging of zebrafish embryos, we previously showed that neural progenitors in the presumptive zebrafish spinal cord are specified prior to and during morphogenetic movements (Xiong et al., 2013, 2018). For example, motor neuron progenitors (pMN cells) are specified at the neural plate stage, in an intermixed pattern with non-pMN cells. Through an unknown mechanism, during tissue morphogenesis the pMN cells sort themselves into a cohesive pMN domain that has sharp boundaries with regions of non-pMN cells (Xiong et al., 2013). These findings make the zebrafish spinal cord an attractive system to study how robust patterning can be achieved despite imprecision in morphogen signaling and extensive cell-cell neighbor exchange during tissue morphogenesis and growth.

In this work, we focused on three neural progenitor domains that form precise patterns during zebrafish spinal cord morphogenesis. We developed cell-based biophysical assays to measure adhesion forces between different neural progenitor pairs, and found that each neural progenitor type exhibits a homotypic binding preference. We identified a three-molecule adhesion code that is unique for each cell type and is the molecular basis for this adhesion preference. The adhesion code is regulated by the same morphogen gradients that instruct cell fate specifications. We propose that this interaction between adhesion-based self-organization and morphogen gradient-based patterning explains how robust pattern formation is possible even in tissues that undergo considerable cell rearrangement during morphogenesis and growth.

## RESULTS

### Neural progenitor cells resolve from mixed positions during specification to form cohesive domains during morphogenesis

We used imaging of live embryos to visualize the dynamics of cell fate specification and formation of neural progenitor domains. We chose to focus on neural progenitor types that can be distinguished by the expression of a single transcription factor. In the spinal cord, the p3, pMN, and p0 progenitors satisfy this criterion as they can be distinguished by the expression of *nkx2.2a*, *olig2*, and *dbx1b*, respectively (Figure 1A) (Briscoe et al., 2000; Gribble et al., 2007; Novitch et al., 2001; Park et al., 2002; Pierani et al., 2001). At the neural tube stage, these three cell types form stereotypical stripe-like domains. To observe the formation of p3, pMN, and p0 domains in live embryos, we used transgenic zebrafish carrying fluorescent reporters of *nkx2.2a*, *olig2*, and *dbx1b* (Kinkhabwala et al., 2011; Kirby et al., 2006; Ng et al., 2005; Shin et al., 2003) (Figure 1B). We use a position metric called V-D position to quantitatively describe the position of each domain along the ventral-to-dorsal axis (Figure S1A-C). A V-D position of 0 corresponds to the ventral-most edge of the spinal cord, while a V-D position of 1 corresponds to the dorsal-most edge of the spinal cord. The *nkx2.2a*-positive p3 domain is the ventral-most domain, directly flanking the Shh-secreting medial floor plate cells, and has an average V-D position of 0.1. The *olig2*-positive pMN domain is adjacent to the p3 domain on its dorsal side, and has an average V-D position of 0.2. The *dbx1b*-positive p0 domain is positioned more dorsally with an average V-D position of 0.55 (Figure 1A-D). The spatial pattern of these reporters in the spinal cord is in excellent agreement with the pattern detected by fluorescent in situ hybridization based on Hybridization Chain Reaction (*in situ* HCR) (Choi et al., 2018) (Figures 1B-D).

**Figure 1.**
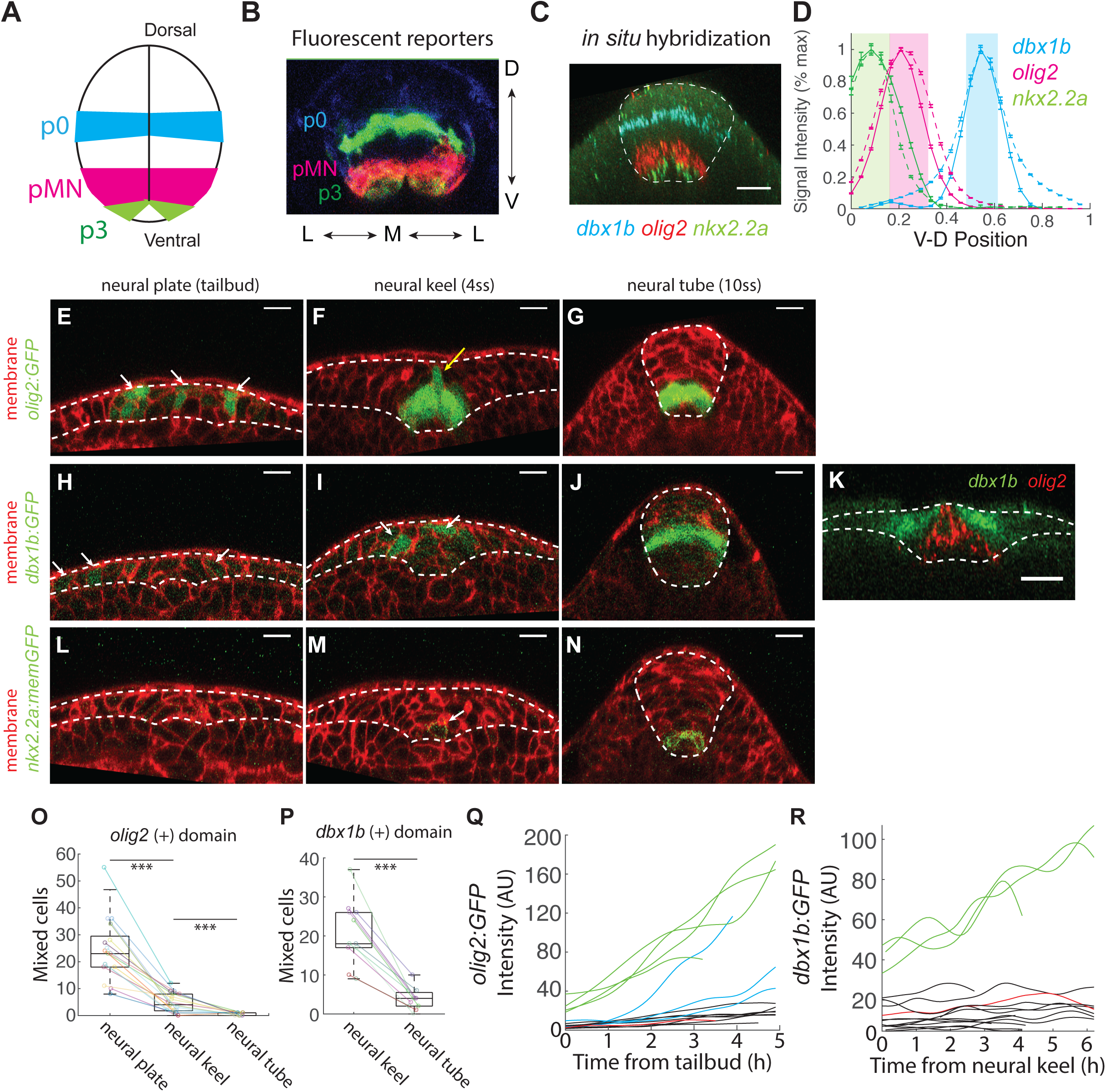
Neural progenitor cells resolve from mixed positions during specification to form cohesive domains during morphogenesis. (A) A cartoon illustration of the stripe patterns of p3, pMN, and p0 domains at the neural tube stage (B) Cross section of the spinal cord of the triple fluorescent reporter fish, *TgBAC(nkx2.2a:memGFP); TgBAC(olig2:dsRed); TgBAC(dbx1b:GFP)*, at 16 somite stage. (C) Cross section of the spinal cord at 11 somite stage, stained with multiplex *in situ* HCR probes against *nkx2.2a* (green), *olig2*(red) and *dbx1b*(blue). The image is the maximal intensity projection over a 3.5 µm wide slab. (D) Average intensity profile of *nkx2.2a*, *olig2*, and *dbx1b* along the ventral-to-dorsal axis of the spinal cord, visualized by either fluorescent reporters (dashed lines) or multiplex *in situ* HCR (solid lines). Error bar represent standard error of the mean. N represents number of embryos analyzed and n represent number of cross sections analyzed. Tg(*nkx2.2a:memGFP*) (N=5, n=505), *TgBAC(olig2:GFP)* (N=11, n= 1462), *TgBAC(dbx1b:GFP)* (N=7, n=867), *in situ* HCR of *nkx2.2a* (N=5, n=505), *olig2* (N=17, n=2277), *dbx1b* (N=12, n=1772). (E-N) Cross section of pMN (E-G), p0 (H-J), p3 (L-N) domains at the neural plate (E,H,L), neural keel (F,I,M) and neural tube (G,J,N) stage. White arrows show fate marker positive cells in discontinuous regions (E,H,I,M). Yellow arrow shows the tip of the wedge-shaped pMN domain at the neural keel stage (F). Scale bars are 20 µm. (K) Fluorescent *in situ* hybridization of *olig2* in *TgBAC(dbx1b:GFP)* fish, showing the p0 cells making contact with the tip of the wedge-shaped pMN domain (O) Quantification of *olig2:GFP* negative cells mixed within the *olig2:GFP* positive domain at the neural plate, neural keel, or neural tube stages across a 265 µm region along the anterior-to-posterior axis. Data are from 17 embryos in 6 experiments. For (O,P), the boxes within the box plot represent 25^th^, 50^th^, and 75^th^ percentile of the data. *** denotes p-value < 0.001 (t-test) (P) Quantification of *dbx1b:GFP* negative cells mixed within the *dbx1b:GFP* positive domain at the neural keel, or neural tube stages across a 265 µm region along the anterior-to-posterior axis. Data are from 11 embryos in 3 experiments. (Q) Timelapse single cell traces of 4 *olig2:GFP* positive (green) and 11 *olig2:GFP* negative cells, starting from the neural plate stage to follow expression of the *olig2:GFP* reporter. The 7 black traces are *olig2:GFP* negative cells that remain *olig2:GFP* negative and sort out from the pMN domain. The 1 red trace is a *olig2:GFP* negative cell that fails to sort from the pMN domain and remains *olig2:GFP* negative. The 3 blue traces are cells that are initially *olig2:GFP* negative, remain in the pMN domain and turn on GFP much later than the neighboring GFP positive cells. The traces are from 4 different embryos in 2 experiments. (R) Single cell traces of 3 *dbx1b:GFP* positive (green) and 12 *dbx1b:GFP* negative cells, starting from the neural keel stage to follow expression of the *dbx1b:GFP* reporter. The 11 black traces are *dbx1b:GFP* negative cells that remain *dbx1b:GFP* negative and sort out from the p0 domain. The 1 red trace is a *dbx1b:GFP* negative cell that fail to sort from the p0 domain and remain mixed. The traces are from 3 different embryos in 2 experiments.

Of these three cell types, the motor neuron progenitors (pMN) show fate-specific gene expression earliest, with clear *olig2:GFP* signal at the neural plate stage (Figure 1E). Fluorescent *in situ* HCR shows that the *olig2* transcript levels are comparable among the neural plate, neural keel, and neural tube stages, validating the early specification of pMN cells observed from the transgenic reporter (Figure S1D). Consistent with our previous findings (Xiong et al., 2013), specified pMN cells in the neural plate exhibit a mixed pattern, with *olig2* positive cells intermixing with *olig2* negative cells (Figure 1E, Movie S1). The mixed pattern is gradually resolved, and a largely cohesive wedge-shaped domain forms by the neural keel stage (Figure 1F, Movie S1 and S2). The wedge shape of the pMN domain is likely due to its compression by neighboring *olig2*-negative cells that are medially converging. The pMN cells at the tip of the wedge (Figure 1F, yellow arrow) typically maintain cohesion with the rest of the pMN domain throughout the neural keel stage, and eventually resolve into a cohesive stripe-like domain at the neural tube stage (Figure 1G, Movie S2).

The p0 cells show fate-specific marker expression shortly after pMN cells. Using fluorescent *in situ* HCR against *dbx1b*, we found that *dbx1b* expression is comparable between the neural keel and neural tube stages, and twice the expression seen at the neural plate stage (Figure 1H-J, S1E). This suggests that the p0 cell identity defined by *dbx1b* expression is established at the neural keel stage. The *dbx1b* positive cells at the left and right side of the neural keel are separated, and mixing of *dbx1b* positive and negative cells is observed locally on both sides of the *dbx1b* domain (Figure 1I and Movie S3). As the tissue continues to converge, the *dbx1b* positive domains from both sides of the neural keel come into contact and merge into a cohesive stripe at the neural tube stage (Figure 1J and Movie S3). We frequently found *dbx1b* positive cells at the neural keel stage in contact with the pMN cells at the tip of the wedge-shaped pMN domain (Figure 1F,I,K).

The p3 cell type expresses its fate marker last. *Nkx2.2a* expression is absent at the neural plate stage, and is first observed at the neural keel stage (Figures 1L,M). *In situ* HCR against *nkx2.2a* transcripts shows monotonic increase from the neural plate stage to the neural tube stage (Figure S1F). At the neural keel stage, we typically observe zero or one p3 cells per cross-section defined by the dorso-ventral and medial-lateral axes (Figure 1M), suggesting that the specified p3 cells are intercalated with non-p3 cells along the anterior-posterior axis. As the tissue enters the neural tube stage, the p3 cells reorganize into two one-cell-wide stripes on both sides of the medial floor plate (Figure 1N and Movie S4). Our time-lapse movies show that the *nkx2.2a*-positive cells remain one to two cell diameters away from the medial floor plate, where Shh is produced, throughout neural tube morphogenesis (Movie S4).

To quantitatively characterize how these neural progenitor domains resolve the initial mixed pattern, we imaged individual embryos from the neural plate to neural tube stage and counted the number of GFP-negative cells mixed within the GFP-positive domain over time in a region corresponding to the 2^nd^ to the 6^th^ somite. On average, 24 *olig2:GFP* negative cells are found in the *olig2:GFP* positive domain at the neural plate stage, but fewer than 5 mixed cells are found at the neural keel stage, suggesting that most mixed cells in the pMN domains move out of this domain between the neural plate to neural keel transition (Figure 1O). For the *dbx1b:GFP* positive domain, we found an average of 20 *dbx1b:GFP* negative cells mixed within the *dbx1b:GFP* positive domain at the neural keel stage, and fewer than 4 mixed cells at the neural tube stage, suggesting that unmixing largely occurs at the neural keel to neural tube transition (Figure 1P).

To visualize how mixing is resolved at a single cell level, we tracked *olig2:GFP* negative cells mixed within the pMN domain, starting at the neural plate stage. Ten out of 11 tracked cells successfully resolved the mixing. Of these, seven cells remained GFP-negative and sorted out from the pMN domain by exchanging locations with their neighbors, while three cells remained within the pMN domain and belatedly turned on the *olig2:GFP* signal (Figure 1Q). Similarly, of thirteen *dbx1b* negative cells mixed within the p0 domain at the neural keel stage, 12 out of 13 cells successfully sorted out from the p0 domain and all twelve remained *dbx1b:GFP* negative (Figure 1R). This suggests that active cell sorting is the primary mechanism that corrects for the initial mixed pattern.

In summary, the pMN and p0 cells are specified early during morphogenesis, in an imprecise pattern that mixes fate marker positive cells with fate marker negative cells. The mixed patterns are resolved during morphogenesis to form cohesive stripe patterns at the neural tube stage. The p3 cells are also specified in a scattered fashion, but as they remain near the source of Shh the movement required to form a cohesive domain is less than that for pMN and p0 cells. All three cell types are specified during tissue morphogenesis, and manage to form cohesive domains that are maintained throughout morphogenetic movement, eventually forming the stereotypic stripe-like pattern.

### Neural progenitor cells exhibit homotypic preference

To test whether different neural progenitor types exhibit differences in their adhesion properties, we first set out to measure adhesion forces between different types of neural progenitor cell contacts. We used the dual pipette aspiration assay to determine the separation forces of different combinations of cell doublets (Figure 2A and Movie S5) (Biro and Maître, 2015; Maître et al., 2012). We dissected the trunk region of neural tube stage (8-10 somite stage) embryos and mechanically dissociated the tissue to obtain single cells, identifying each of the three neural progenitor types based on their expression of fluorescent markers (Figure 1). We measured the adhesion forces of six different combinations of cell doublets (Figure 2B), including three different homotypic contacts (contacts between cells of the same type) and three different heterotypic contacts (contacts between cells of different types). The adhesion at the pMN-pMN homotypic contact is strongest (7.6 ± 3.6 nN), and is significantly greater than the pMN-p3 heterotypic contact (2.5 ± 2.2 nN). The p3-p3 homotypic contact (4.0 ± 2.7 nN) is also significantly stronger than the pMN-p3 heterotypic contact, even though the p3 cells do not appear to face the challenge of sorting over a large distance (Figure 2B). Similarly, the average adhesion forces at the homotypic contacts between two pMN cells (7.6 ± 3.6 nN) and two p0 cells (7.2 ± 6.3 nN) are also greater than the pMN-p0 heterotypic contacts (4.4 ± 3.3 nN) (Figure 2B). Thus, each of the three neural progenitor types exhibit homotypic preference, a term we use to describe the phenomenon that cells selectively stabilize homotypic contacts over heterotypic contacts.

**Figure 2.**
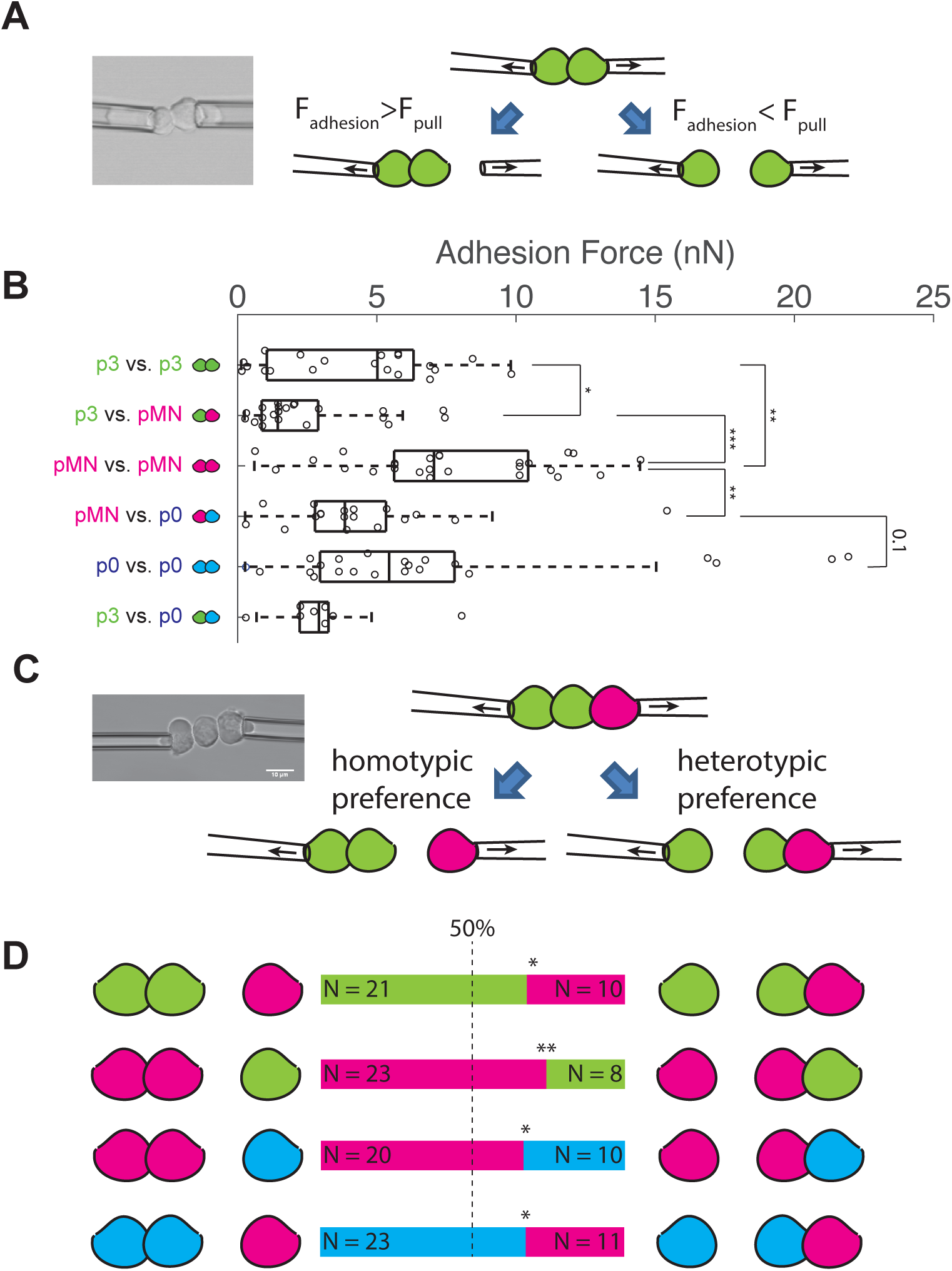
Neural progenitor cells exhibit homotypic preference. (A) A cartoon illustration of the dual pipette aspiration assay. (B) The adhesion force measured from 6 types of cell doublets. The boxes within the box plot represent 25^th^, 50^th^, and 75^th^ percentile of the data. N denotes the number of independent experiments and n denotes the number of measurements. p3-p3 (N=9, n= 37), p3-pMN (N=5, n=22), pMN-pMN (N=6, n=26), pMN-p0 (N= 3, n=18), p0-p0 (N=5, n=22), p3-p0 (N=2, n=8). * and ** represent p-values <0.05 and <0.01, respectively (t-test) (C) A cartoon illustration of the triplet assay. Scale bar of the inset is 10 µm. (D) Triplet assays to observe homotypic preference. Green, magenta, and blue represent p3, pMN and p0 cells. N denotes the number of independent experiments and n denotes the number of measurements. p3-p3-pMN (N=7, n=31), pMN-pMN-p3 (N=5, n=31), pMN-pMN-p0 (N=6, n=30), p0-p0-pMN (N=6, n=34). * and ** represent p-values <0.05 and <0.01, respectively (binomial test)

To enable more sensitive comparison of preferences between distinct contact types, we further developed a novel triplet competition assay (Figure 2C). The triplet is composed of two cells of the same type and one cell of a different type, forming one homotypic contact and one heterotypic contact. This assay mimics the challenge faced by the cells when they are pulled by neighboring cells towards different directions *in vivo*. We always started with three isolated cells, and assembled them into a triplet configuration by simultaneously establishing both contacts (Movie S6). The three cells were allowed to form contacts for at least 3 minutes and then pulled apart by stage-motor controlled micropipettes. As the micropipettes pull at the cells on either side of the triplet, the triplet will break apart. The middle cell will stay with either the right cell or the left cell, generating a direct competition between the homotypic and heterotypic contact. All three neural progenitor cell types showed clear homotypic preference in this assay, with pMN-pMN and p3-p3 homotypic contacts winning over pMN-p3 heterotypic contact, and pMN-pMN and p0-p0 homotypic contacts winning over pMN-p0 heterotypic contact by a ratio of approximately 2:1 (Figure 2D). Thus, both the doublet and triplet assays revealed a preference for p3, pMN, and p0 cells to adhere more strongly to other cells of the same type.

### Transcriptomic and mutational analyses reveal three key adhesion molecules for the patterning of neural progenitors

The homotypic preference we observed in the three neural progenitor types suggest that each cell type may utilize distinct adhesion machineries to achieve adhesion specificity. To identify the molecular mechanism underlying this specificity, we isolated p3, pMN, and p0 cells by fluorescence-assisted cell sorting of single cells dissociated from embryos with fluorescent fate reporters (Figure 1), and performed RNA-seq to identify the gene expression profile of each cell type (Figure 3A,B). The cadherin/protocadherin families of adhesion molecules have frequently been observed to mediate homotypic interactions (Frank and Kemler, 2002; Nose et al., 1988; Steinberg and Takeichi, 1994). 13 of the genes encoding members of this family have expression levels above our abundance threshold in at least one cell type, and 7 of these were expressed differentially by more than two-fold between different cell types (Table S1). To identify which of these molecules are relevant to neural progenitor patterning, we used CRISPR-Cas9-mediated genome editing to knock out each of the 13 genes in our fish lines that carry fluorescent reporters for the three neural progenitor domains, and imaged the embryos injected with the guide RNA-Cas9 mix to look for patterning phenotypes in the neural tube at the 10 somite stage (Gagnon et al., 2014). We were able to assess spinal cord patterning in all the CRISPR mutants except for E-cadherin (*cdh1*) CRISPR mutants, which exhibited severe gastrulation defects. N-cadherin (*cdh2*), cadherin 11 (*cdh11*), and protocadherin 19 (*pcdh19*) stood out as genes with significant loss-of-function phenotypes (Figure 3C-E). Knocking out any one of these three genes affected the patterning of the p3 and pMN domains, while loss of *cdh2* and *pcdh19* also affected p0 domain patterning. Of the 12 CRISPR mutants we were able to study, only the *cdh2* CRISPR mutants exhibited obvious embryonic morphological defects. The other 11 CRISPR mutants do not have gross morphological defects in the spinal cord at the 10 somite stage.

**Figure 3.**
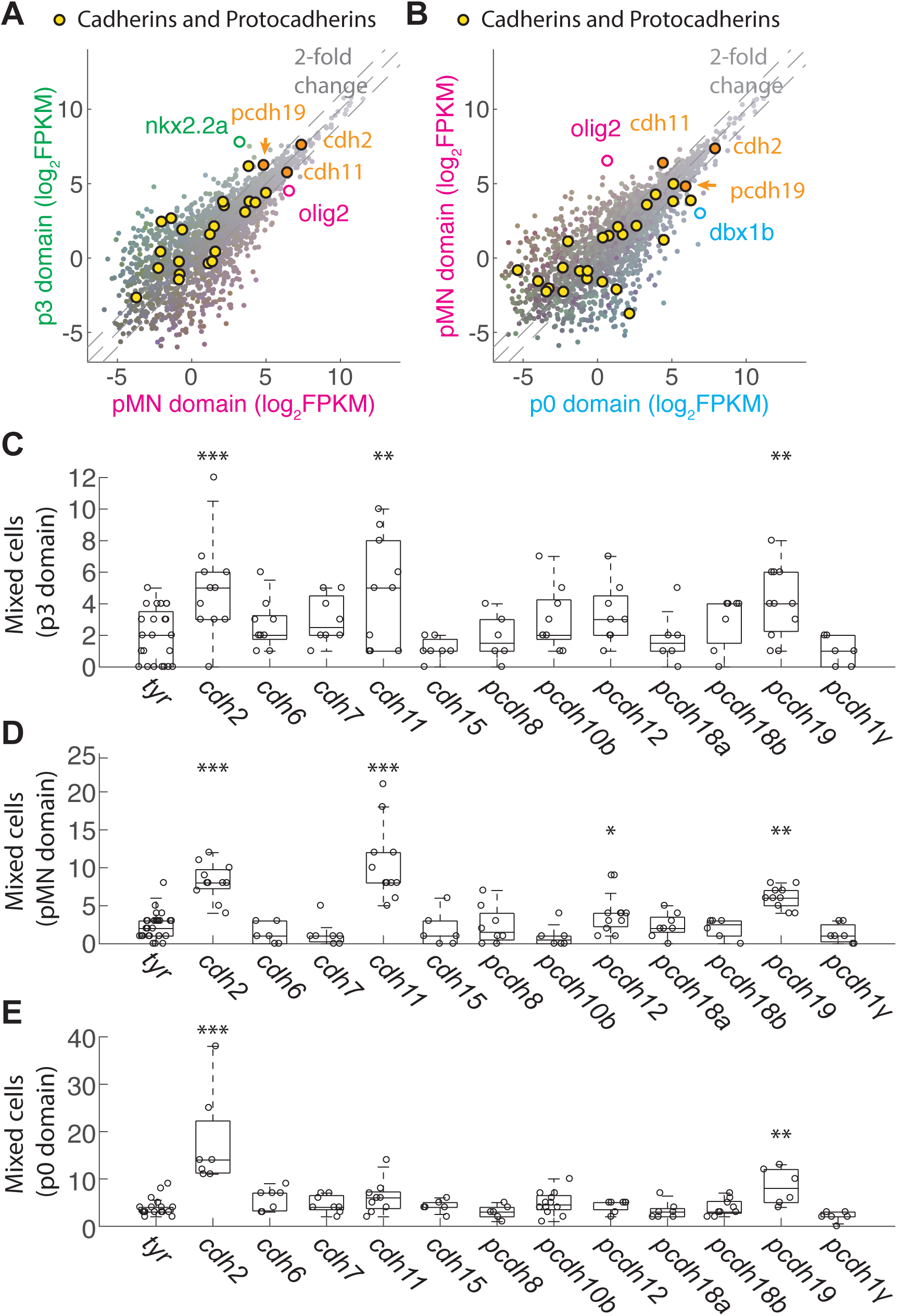
Transcriptomic and mutational analyses reveal three key adhesion molecules for the patterning of neural progenitors. (A-B) RNAseq of pMN cells versus p3 cells (A), or p0 cells versus pMN cells (B). The genes belonging to the cadherin or protocadherin family are labeled with yellow circles, except for *cdh2*, *cdh11* and *pcdh19*, which are labeled with orange circles. In (A), points to the left of the gray dashed lines represent genes enriched in the p3 domain by more than 2-fold, such as the fate marker for p3 cells, *nkx2.2a*. Points to the right of the gray dashed lines represent genes enriched in the pMN domain by more than 2-fold, such as the fate marker for pMN cells, *olig2*. In (B), points to the left of the gray dashed lines represent genes enriched in the pMN domain by more than 2-fold, such as the fate marker for pMN cells, *olig2*. Points to the right of the gray dashed lines represent genes enriched in the p0 domain by more than 2-fold, such as the fate marker for p0 cells, *dbx1b*. (C-E) Results of CRISPR genetic screen in the p3 (C), pMN(D), or p0 (E) domains. In (C), transgenic embryos *TgBAC(nkx2.2a:memGFP);TgBAC(olig2:dsRed)* were used. In (D), transgenic embryos *TgBAC(olig2:GFP); Tg(actb2:memCherry2)* were used. And in (E), transgenic embryos *TgBAC(dbx1b:GFP); Tg(actb2:memCherry2)* were used. CRISPR guide RNA against tyrosinase was used as control. The boxes within the box blot represent 25^th^, 50^th^, and 75^th^ percentile of the data. *, **, and *** represent p-value of <0.05, <0.01, <0.001, respectively (t-test). The following list represents the number of independent experiments and the number of embryos used in each condition. (C) tyr (5,23), *cdh2* (2,11), cdh6 (1,9), cdh7 (2,8), *cdh11* (1,10), cdh15 (1,7), pcdh8 (1,6), pcdh10b (1,9), pcdh12 (1,8), pcdh18a (1,6), pcdh18b (1,7), *pcdh19* (1,11), pcdh1γ (1,6). (D) tyr (5,25), *cdh2* (2,11), cdh6 (1,6), cdh7 (1,7), *cdh11* (1,11), cdh15 (1,6), pcdh8 (1,8), pcdh10b (1,6), pcdh12 (1,11), pcdh18a (1,8), pcdh18b (1,6), *pcdh19* (1,10), pcdh1γ (1,7) (E) tyr (5,17), *cdh2* (2,7), cdh6 (2,7), cdh7 (3,8), *cdh11* (1,9), cdh15 (1,6), pcdh8 (2,6), pcdh10b (4,12), pcdh12 (1,7), pcdh18a (3,7), pcdh18b (2,9), *pcdh19* (2,6), pcdh1γ (1,6).

### Differential expression of *cdh2*, *cdh11*, and *pcdh19* forms a unique combinatorial adhesion code for each cell type

We next sought to determine the spatiotemporal patterns of expression of *cdh2*, *cdh11* and *pcdh19* during neural progenitor patterning. Cdh2 is the most ubiquitous cadherin throughout the nervous system. Its expression starts in the late gastrula and persists through the transition from the neural plate to the neural tube (Harrington et al., 2007; Lele et al., 2002) Our transcriptomic study shows that *cdh2* is the most abundant cadherin in p3, pMN, and p0 cells (Figure 3A,B). Using a fluorescent reporter, *TgBAC(cdh2:cdh2-mCherry)* (Colak-Champollion et al., 2019), we found that the Cdh2 protein exhibits a linear gradient along the V-D axis of the neural tube that increases by 2-fold from the V-D position of 0 to the 0.8 position (Figure 4A-C) before decreasing towards the dorsal-most part of the neural tube. This ventral-to-dorsal pattern of Cdh2 can be observed throughout the entire spinal cord (Figure 4A). By superimposing the V-D position of the p3, pMN, and p0 cells on the Cdh2 gradient, we estimated the Cdh2 level in the locations of the different domains. We estimate that if we set the level of Cdh2 at the p0 domain as 1 in arbitrary units, then the p3 domain averages a level of 0.6, and pMN domain averages a level of 0.7-0.8 (Figure 4C,N). Similar spatial profiles of Cdh2 are also observed by antibody staining of sectioned neural tube (Figure S3A,B) and in a different fluorescent reporter fish of *cdh2*, *TgBAC(cdh2:cdh2-tFT)* (data not shown) (Revenu et al., 2014). Interestingly, this pattern does not appear in *in situ* HCR for *cdh2* mRNA in neural tube sections, where we observed constant levels of *cdh2* mRNA between V-D position 0 to 0.8 and reduced *cdh2* mRNA levels only at the dorsal-most part of the neural tube (Figure S3C,D). This is consistent with our transcriptomic study, which showed similar levels of *cdh2* mRNA in the p3, pMN, and p0 domains (Figure 3A,B), and indicates that the Cdh2 protein gradient may be regulated post-transcriptionally.

**Figure 4.**
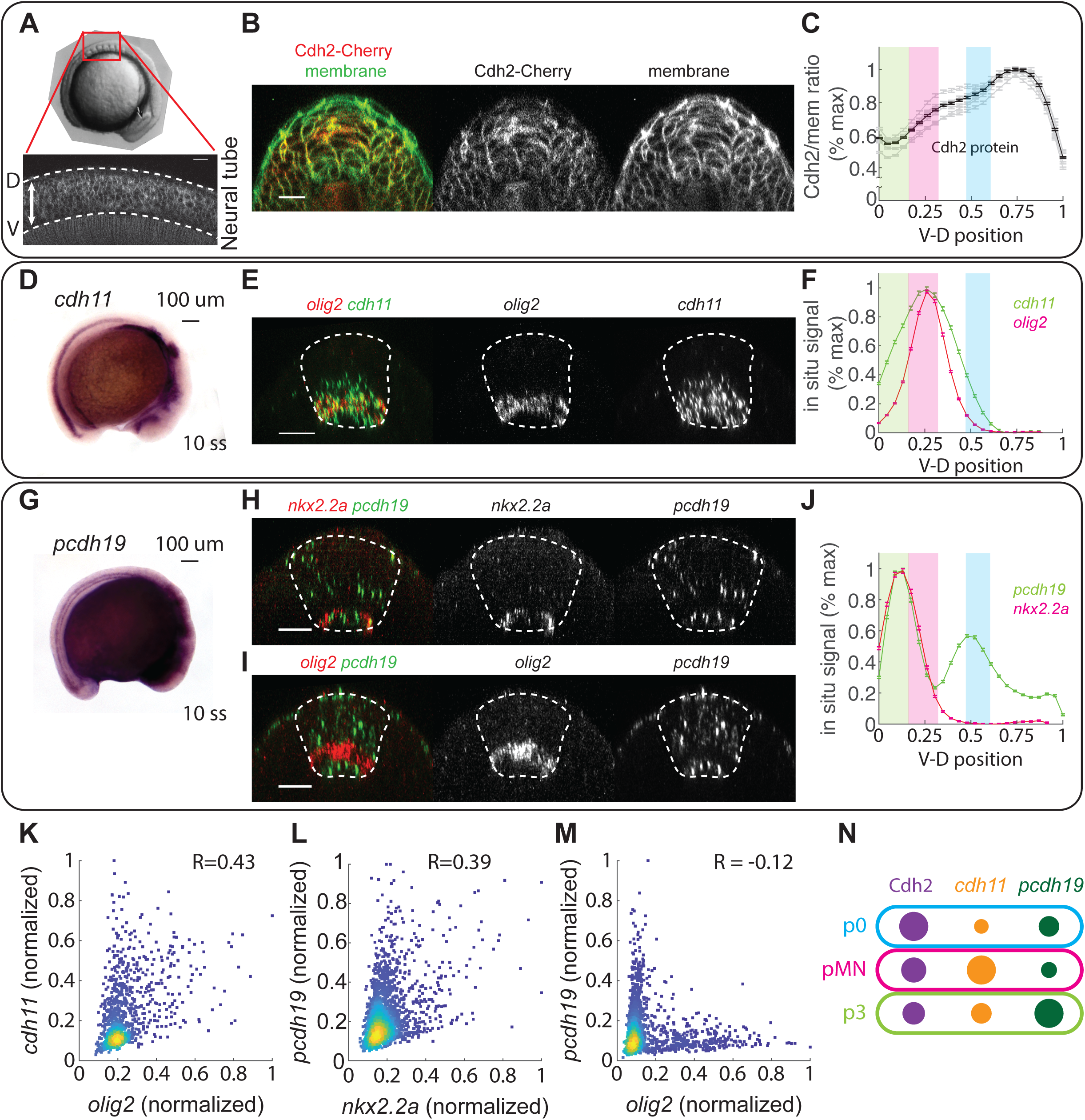
Differential expression of *cdh2*, *cdh11*, and *pcdh19* forms a unique combinatorial adhesion code for each cell type. (A) Lateral view of a 10 somite stage embryo, showing the Cdh2 gradient in the neural tube of an transgenic embryo *TgBAC(cdh2:cdh2-Cherry)* (B) Cross section of a confocal z-stack of a neural tube from a 10-somite stage transgenic embryo with *TgBAC(cdh2:cdh2-Cherry); Tg(actb2:membrane-Citrine)*. Scale bar is 20 µm. (C) Ventral-to-dorsal Intensity profile of *cdh2*-Cherry, normalized by the membrane-Citrine signal. Gray lines in the background are average intensity profiles from different embryos. The shaded regions of green, magenta, and blue in (C,F,J) represent the positions of p3, pMN, and p0 domains, and were identical to Figure 1D. Data are from 606 line traces in 6 embryos. Error bars are standard error of the mean. (D, G) Representative images from whole mount in situ hybridization of *cdh11* (D) and *pcdh19* (G) at the neural tube stage (10 somite stage) (E) Cross-section of a 10-somite stage embryo with multiplex *in situ* HCR staining with probes against *olig2* and *cdh11*, image is the maximal intensity projection over 1.79 µm wide slices. Scale bar is 20 µm. (F) Ventral-to-dorsal intensity profile of *olig2* and *cdh11* based on multiplex *in situ* HCR staining. An average of 3212 intensity profiles in 31 embryos from 4 independent experiments is shown. Error bars are standard error of the mean. (H) Cross-section of a 10-somite stage embryo with *in situ* HCR staining with *nkx2.2a* and *pcdh19*, image is the maximal intensity projection over 4.87 µm wide slices. Scale bar is 20 µm. (I) Cross-section of a 10-somite stage embryo with *in situ* HCR staining with *olig2* and *pcdh19*, image is the maximal intensity projection over 2.66 um wide slices. Scale bar is 20 µm. (J) Ventral-to-dorsal intensity profile of *nkx2.2a* and *pcdh19* based on multiplex *in situ* HCR staining. An average of 2121 intensity profiles in 20 embryos from 4 independent experiments is shown. Error bars are standard error of the mean. (K-M) Single cell co-expression pattern of *olig2*-*cdh11* (K), *nkx2.2a* – *pcdh19* (L), and *olig2* – *pcdh19* (M) quantified from embryos stained with multiplex *in situ* HCR. The following number of embryos and cells are used in each panel: (K) (3,1130) (L) (6, 2025) (M) (4, 1688) (N) A cartoon diagram showing the expression of *cdh2*, *cdh11*, and *pcdh19* in the three neural progenitor types. The area of each dot represents the relative abundance of each adhesion molecule, defined by the *cdh2*-cherry level in (C) and the fluorescent intensity of *in situ* HCR for *cdh11* (F) and *pcdh19* (I). The intensity value of each cell type is estimated from the center of the shaded region, where the fate marker expression is at its maximum.

The expression of *cdh11* appears as one stripe along the entire spinal cord (Figure 4D), largely overlapping with the *olig2* positive domain but with a wider distribution along the V-D axis (Figure 4E,F). Quantification of the *cdh11* fluorescent *in situ* signal agrees with our transcriptome analysis: setting the *cdh11* expression level in the pMN domain at 1, the p3 domain shows a level of ∼0.6 and the p0 domain shows a level of ∼0.25 (Figure 4F,N).

Finally, *pcdh19* is expressed as two stripes along the entire spinal cord (Figure 4G). Fluorescent *in situ* analysis shows that the ventral stripe of *pcdh19* expression corresponds to the p3 domain and the medial floor plate (Figure 4H), while the dorsal stripe is positioned dorsal to the pMN domain (Figure 3I). The two *pcdh19* expression domains flank the pMN domain and expression of *pcdh19* and *olig2* appear to be mutually exclusive (Figure 4I,J). We estimate that if we set the level of *pcdh19* in the p3 domain to 1, the pMN domain shows a level of ∼0.3 and the p0 domain shows a level of ∼0.5 (Figure 4J,N).

To further quantify the co-expression of cell fate markers and the adhesion molecules at single cell resolution, we made use of transgenic embryos ubiquitously expressing membrane markers to outline individual cells. The membrane signal is well-preserved throughout multiplex *in situ* HCR (Figure S3E). We manually labelled the center of individual cells, and quantified the average fluorescent intensity of each probe in a cube of 3.5µm on each side within each cell. This single cell co-expression analysis revealed a high correlation between *cdh11* and *olig2* expression (Figure 4K). Almost all *olig2*-positive cells are *cdh11*-positive, while some *cdh11*-positive cells are *olig2*-negative, consistent with the fact that the distribution of *cdh11* expression is wider than the *olig2* domain (Figure 4F). Similarly, *pcdh19* and *nkx2.2a* expression are highly correlated. Almost all *nkx2.2a*-positive cells are *pcdh19*-positive, but a significant portion of *pcdh19*-positive cells are *nkx2.2a*-negative. These cells are mostly positioned within the dorsal-stripe of *pcdh19* expression (Figure 4L). On the other hand, *pcdh19* and *olig2* expression appear to be mutually exclusive even at the single-cell level (Figure 4M). Together, expression of *cdh2*, *cdh11*, and *pcdh19* constitutes a three-molecule adhesion code that is unique to each of the three cell types (Figure 4N).

Importantly, all three adhesion molecules are expressed at the neural plate stage with a pattern consistent with the later neural tube stage. At the neural plate stage, *cdh2* is expressed ubiquitously (Figure S3F), and the Cdh2 protein gradient emerges as the tissue converges to form the neural keel (Figure S3G). *Cdh11* is expressed as a stripe in the medial side of the neural plate, overlapping with the *olig2* expression domain (Figure S3H). *Pcdh19* is expressed in one medial and two lateral stripes that are mutually exclusive with the *olig2* domain (Figure S3I). This suggests that differential expression patterns of *cdh2*, *cdh11*, and *pcdh19* are present throughout neural tube morphogenesis, making these genes plausible candidates for regulators of the formation and maintenance of neural progenitor patterns.

### Cdh11 and Pcdh19 mediate the homotypic preference between pMN and p3 cells

To determine whether these cell adhesion molecules mediate homotypic preference in each of the three neural progenitor types, we first focused on the two ventral cell types, pMN and p3 cells. Among the three neural progenitor types, *cdh11* is enriched in the pMN cells (Figure 4F,N). Cdh11 belongs to type II cadherins, a family of cadherins that structural analyses suggest are unable to bind in trans with type I cadherins (e.g. Cdh2) (Patel et al., 2006). Cdh11-expressing cells and Cdh2-expressing cells were also shown to segregate in culture (Kimura et al., 1995). This suggests that Cdh11 and Cdh2 could mediate mutually exclusive homophilic adhesion, and indicates that Cdh11 is a plausible candidate to mediate pMN-specific homotypic adhesion in a tissue that is otherwise dominated by Cdh2-based adhesion. To determine whether Cdh11 mediates the homotypic preference of pMN cells, we performed the triplet assay on cells isolated from *cdh11* knocked-down embryos. Without Cdh11, pMN cells no longer exhibited homotypic preference against p3 cells (Figure 5A). Adhesion force measurements of cell doublets further showed that loss of Cdh11 specifically lowers pMN-pMN homotypic adhesion without affecting pMN-p3 heterotypic adhesion (Figure 5B). Therefore, the pMN-specific expression of Cdh11 increases pMN-pMN homotypic adhesion, leading to homotypic preference of pMN cells against p3 cells.

**Figure 5.**
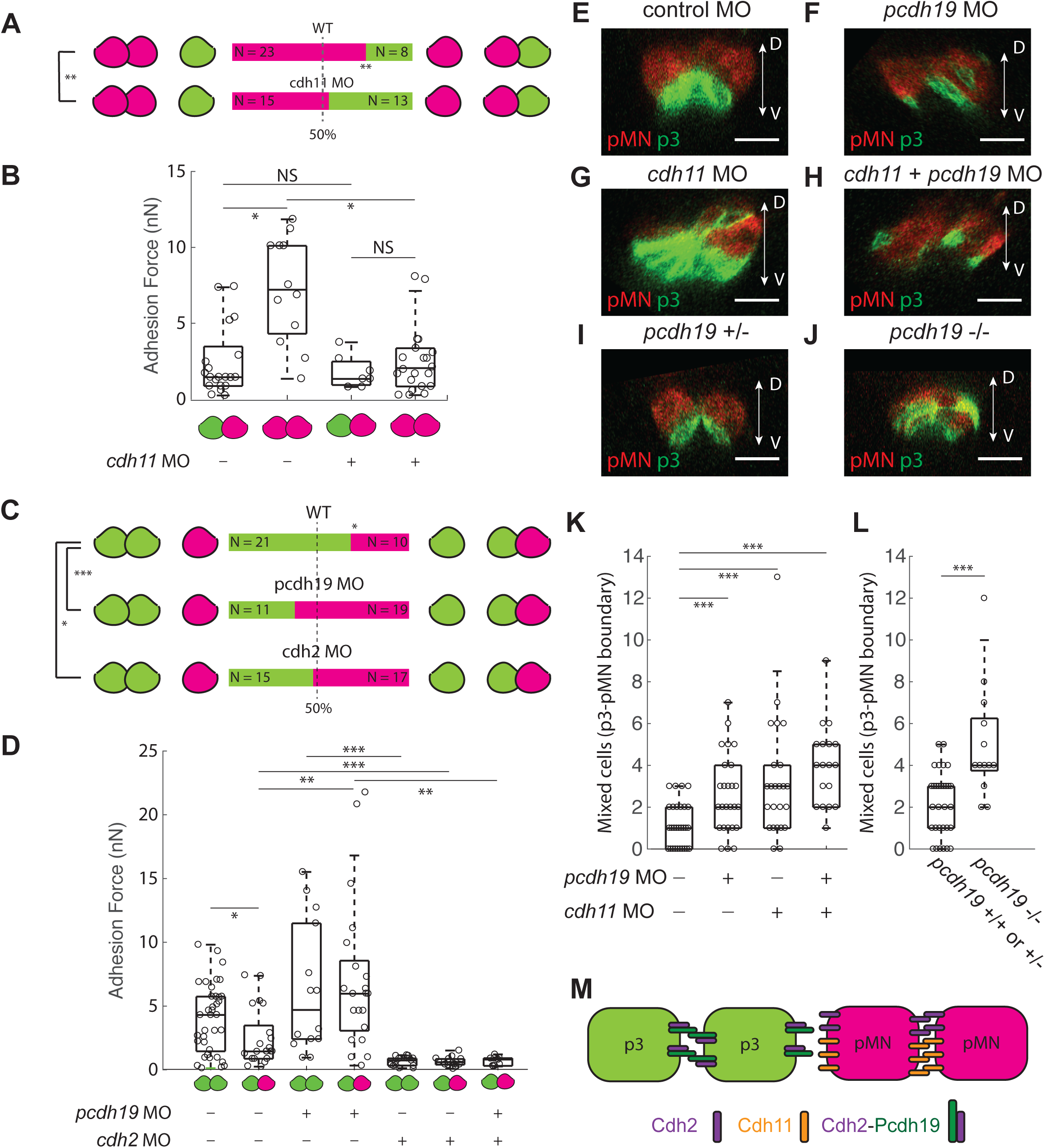
Cdh11 and Pcdh19 mediate the homotypic preference between pMN and p3 cells. (A) Results of the pMN-pMN-p3 triplet assay with and without *cdh11*. For each condition, N denotes number of separate experiments and n denotes number of measurements. Control embryos (N=5, n=31), *cdh11* morpholino-injected embryos (N=3, n=28). ** represent p-value <0.01 (binomial test). (B) Adhesion force of pMN-pMN homotypic adhesion and pMN-p3 heterotypic adhesion, with and without *cdh11*. pMN-p3 in control embryos (N=3, n=21), pMN-pMN in control embryos (N=4, n=12) pMN-p3 in *cdh11* morphant embryos (N=1, n=7), pMN-pMN in *cdh11* morphant embryos (N=6, n=21). For (B,D), the boxes within the box plot represent 25^th^, 50^th^, and 75^th^ percentile of the data. *, **, and *** represent p-value <0.05, <0.01, and <0.001, respectively (t-test). (C) Results of the p3-p3-pMN triplet assay with and without *pcdh19* or *cdh2*. Control (N=7, n=31), *pcdh19* MO (N=5, n=30), *cdh2* MO (N=6, n=32). * and *** represent p-value <0.05 and <0.001, respectively (binomial test). (D) Adhesion force of p3-p3 homotypic adhesion and pMN-p3 heterotypic adhesion, with and without *pcdh19* and *cdh2*. p3-p3 in control embryos (N=9, n= 37), p3-pMN in control embryos (N=5, n=22), p3-p3 in *pcdh19* morphant embryos (N=4, n=14), p3-pMN in *pcdh19* morphant embryos (N=6, n=22), p3-p3 in *cdh2* morphant embryos (N=4, n=15), p3-pMN in *cdh2* morphant embryos (N=3, N=16), p3-pMN in *pcdh19*, *cdh2* double morphant embryos (N=1, n=9). (E-H) p3-pMN boundary at the neural tube stage, visualized by TgBAC(*nkx2.2a:memGFP);TgBAC(olig2:dsRed)*. Embryos were injected with (E) 2.1 ng of standard control morpholino, (F) 4.2 ng of *pcdh19* morpholino, (G) 2.1 ng of *cdh11* morpholino, or (H) *pcdh19* and *cdh11* morpholino (2.1 ng each). Scale bars are 20 µm. Images were oriented with dorsal side of the neural tube at the top. (I-J) p3-pMN boundary at the neural tube stage, visualized by *TgBAC(nkx2.2a:memGFP);TgBAC(olig2:dsRed)* in a *pcdh19* heterozygote (I) or *pcdh19* mutant (J) background. Scale bars are 20 µm. Images were oriented with dorsal side of the neural tube at the top. (K) Quantification of mixed cells at the p3-pMN boundary in embryos injected with standard control morpholino (N=4, n=29), *pcdh19* MO (N=4, n=26), *cdh11* MO (N=2, n=24), or *pcdh19* + *cdh11* MO (N=2, n=18). For (K,L), *** represents p-value < 0.001 (t-test) (L) Quantification of mixed cells at the p3-pMN boundary in *pcdh19* mutant embryos (N=2, n=31) versus its wildtype or heterozygote siblings (N=2, n=13) (M) A cartoon diagram summarizing the working model for homotypic preference between the p3 and pMN cells. The p3 cells utilize the Pcdh19-Cdh2 complex while the pMN cells utilize Cdh2 and Cdh11 to achieve homotypic preference.

We hypothesized that Pcdh19 is responsible for the homotypic preference of p3 cells against pMN cells, as it is specifically enriched in the p3 cells over pMN cells (Figure 4H-J). Elegant *in vitro* studies have shown that Pcdh19 can only generate very weak adhesion on its own (Cooper et al., 2016). However, Pcdh19 can form a complex with Cdh2 to generate strong homophilic adhesion in *trans* using the extracellular domains of Pcdh19, possibly due to a conformational change in Pcdh19 mediated by Cdh2 binding (Emond et al., 2011). Importantly, the Cdh2-Pcdh19 complex does not form adhesions with other Cdh2 molecules in *trans*, suggesting that the Pcdh19-Cdh2 complex and Cdh2 alone have mutually exclusive adhesion modes (Emond et al., 2011). Since Cdh2 is expressed ubiquitously throughout the neural tube, the presence or absence of Pcdh19 in different cell types might thus generate two distinct modes of cell adhesion. As p3 cells express very high levels of *pcdh19* mRNA, while pMN cells express very low levels (Figure 4N), we examined whether Pcdh19 mediates the homotypic preference of the Pcdh19-positive p3 cells against the Pcdh19-negative pMN cells in the triplet assay. Loss of either Pcdh19 or Cdh2 removed the preference for p3-p3 binding versus p3-pMN binding in our triplet assay (Figure 5C), consistent with the idea that a complex of Pcdh19 and Cdh2 is required for p3 homotypic preference.

To determine how changes in Pcdh19 and Cdh2 levels affect p3-p3 homotypic adhesion and p3-pMN heterotypic adhesion, we measured adhesion force in our doublet assay. Interestingly, loss of *pcdh19* does not change the p3-p3 homotypic adhesion force significantly; instead, the predominant effect of the loss of Pcdh19 is an increase of the p3-pMN heterotypic adhesion to a level that is comparable with the p3-p3 homotypic adhesion (Figure 5D). Based on the reported behavior of the Pcdh19-Cdh2 complex, we speculated that p3-pMN heterotypic adhesion is weak because the Pcdh19-Cdh2 complex formed in p3 cells cannot bind to the Cdh2 proteins on pMN cells. Removal of *pcdh19* in p3 cells would then allow Cdh2-based adhesion both between p3 cells and between p3 and pMN cells. Since Cdh2 levels are similar between the two cell types (Figure 4C,N) heterotypic adhesion would then be essentially identical to homotypic adhesion. We therefore predicted that the increased p3-pMN heterotypic adhesion in embryos without *pcdh19* is Cdh2-dependent and should vanish if Cdh2 is also lost. Indeed, when we knocked down *cdh2* and *pcdh19* simultaneously, the p3-pMN adhesion force was reduced to a negligible level (Figure 5D). *Cdh2* knockdown alone also reduces p3-pMN adhesion to almost nothing. Together, our data suggest that Pcdh19 does not directly function to strengthen p3-p3 homotypic adhesion, but instead generates a distinct adhesion mode that lowers p3-pMN heterotypic adhesion.

In our CRISPR screen, loss of *cdh2*, *cdh11*, or *pcdh19* each disrupted patterning of the p3 and pMN domains (Figure 3C,D). We also knocked down *pcdh19*, *cdh11*, or both using morpholinos to confirm these results using an alternative perturbation. Consistent with our CRISPR results, we observed an increase in cell mixing across the p3-pMN boundary in single or double morphants of *cdh11* and *pcdh19* (Figure 5E-H,K). We also crossed the fish line carrying fluorescent reporters of the p3 and pMN domains with a *pcdh19* mutant generated by TALEN-based gene editing (Cooper et al., 2015). This allowed us to compare the number of mixed cells across the p3-pMN boundary in homozygote *pcdh19* mutants versus their wildtype or heterozygote siblings, and confirmed the patterning defect seen in our *pcdh19* CRISPR mutant (Figure 5I,J,L). Together, the CRISPant, morphant, and mutant results suggest that *pcdh19* and *cdh11* are important mediators of the formation of a sharp boundary between the p3 and pMN domains *in vivo*. Our results provide strong support for the conclusion that differential expression of *cdh11* and *pcdh19* not only mediates the homotypic preference of p3 and pMN cells *ex vivo*, but also maintains the boundary between the p3 and pMN domains *in vivo* (Figure 5M).

### Cdh11 and Cdh2 mediate the homotypic preference between pMN and p0 cells

We next aimed to identify the molecules responsible for the homotypic preference of pMN and p0 cells. As is the case for the homotypic preference of pMN cells against p3 cells, Cdh11 also mediates the homotypic preference of pMN cells against p0 cells. Without Cdh11, pMN cells lost homotypic preference against p0 cells, and pMN-p0 heterotypic adhesion became favored over the pMN-pMN homotypic adhesion in the triplet assay (Figure 6A). In the adhesion force measurement, loss of Cdh11 also specifically reduced pMN-pMN homotypic adhesion without affecting pMN-p0 heterotypic adhesion (Figure 6B). Since Cdh2 is abundantly expressed in both pMN and p0 cells, we also performed the pMN-pMN-p0 triplet assay in the absence of Cdh2. Remarkably, pMN-pMN homotypic adhesion became even more favored over pMN-p0 heterotypic adhesion (Figure 6A). This is again consistent with the adhesion force measured from cell doublets. Without Cdh2, p0-pMN heterotypic adhesion became negligible (<1 nN), while the pMN-pMN homotypic adhesion still maintained about 30% of its wildtype level (2-3 nN). We next tested whether the remaining adhesion in Cdh2-deficient pMN cells is provided by Cdh11. Indeed, when *cdh2* and *cdh11* are knocked down simultaneously, pMN-pMN homotypic adhesion decreased to a negligible level (<1nN). Together, these results suggest that Cdh2 and Cdh11 based adhesion are roughly equally responsible for pMN-pMN homotypic adhesion, while pMN-p0 heterotypic interactions are almost entirely mediated by Cdh2 based adhesion. The pMN homotypic preference against p0 cells is driven by the additional Cdh11-mediated pMN-pMN adhesion.

**Figure 6.**
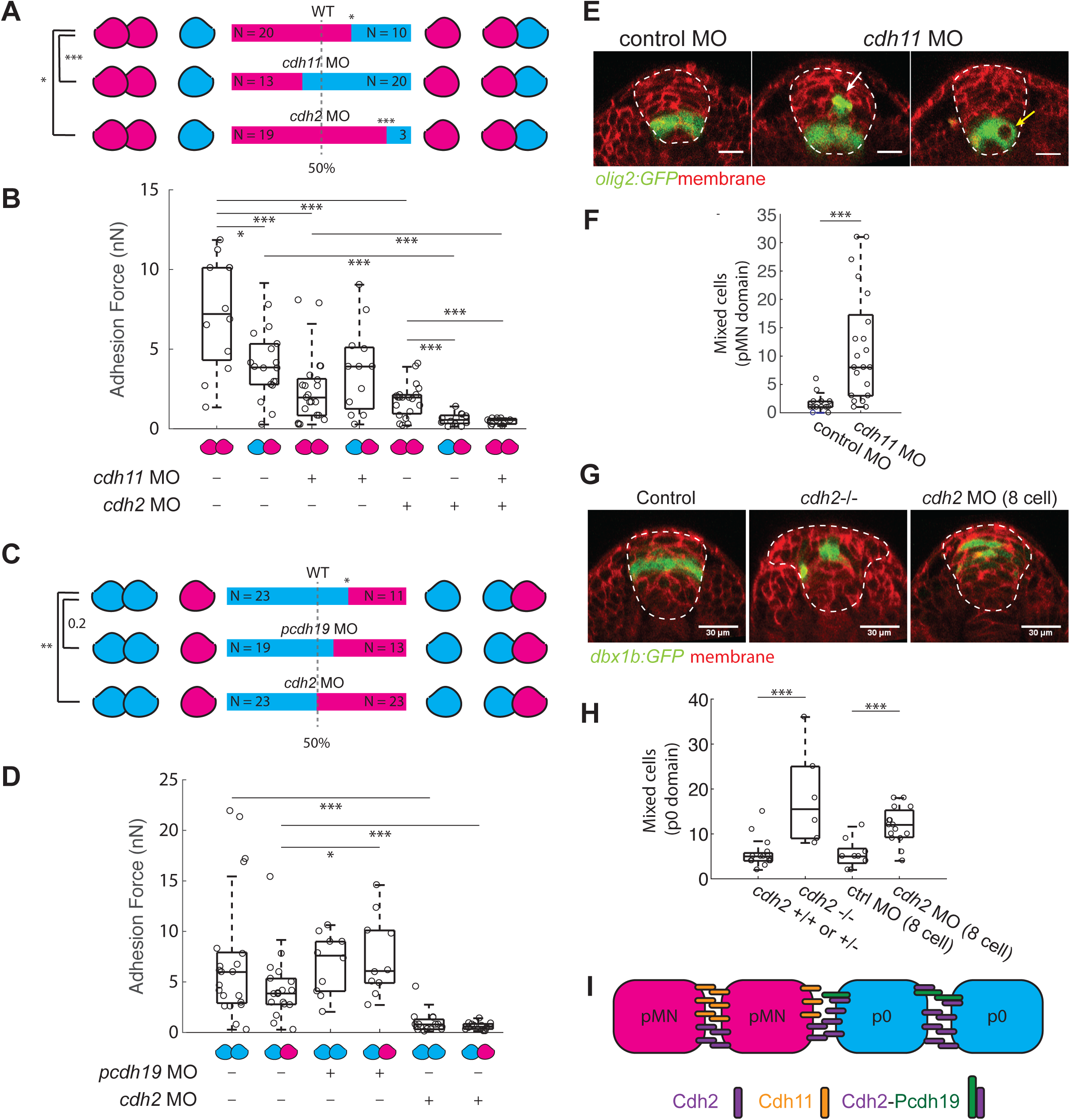
Cdh11 and Cdh2 mediate the homotypic preference between pMN and p0 cells. (A) Results of the pMN-pMN-p0 triplet assay with and without *cdh11* or *cdh2*. Control (N=6, n=30), *cdh11* morphant (N=5, n=33), *cdh2* morphant (N=5, n=22). For (A,C), *,** and *** represent p-value <0.05, <0.01, and <0.001 (binomial test). (B) Adhesion force of pMN-pMN homotypic adhesion and pMN-p0 heterotypic adhesion, with and without *cdh11* or *cdh2*. pMN-pMN in standard control morpholino-injected embryos (N=4, n=12), pMN-p0 in standard control morpholino-injected embryos (N=4, n=18), pMN-pMN in *cdh11* morphant embryos (N=6, n=21), pMN-p0 in *cdh11* morphant embryos (N=3, n=12), pMN-pMN in *cdh2* morphant embryos (N=5, n=20), pMN-p0 in *cdh2* morphant embryos (N=3, n=12), pMN-pMN in *cdh2* and *cdh11* double morphant embryos (N=1, n=11). For (B,D), the boxes within the box plot represent 25^th^, 50^th^, and 75^th^ percentile of the data. * and *** represent p-value < 0.05 and <0.001 (t-test). (C) Results of the p0-p0-pMN triplet assay with and without *pcdh19* or *cdh2*. Control embryos (N=6, n=34), *pcdh19* morphant embryos (N=6, n=32), *cdh2* morphant embryos (N=7, n=46). (D) Adhesion force of p0-p0 homotypic adhesion and p0-pMN heterotypic adhesion, with and without Pcdh19 and Cdh2. p0-p0 in control embryos (N=5, n=22), p0-pMN in control embryos (N=3, n=18), p0-p0 in *pcdh19* morphant embryos (N=4, n=10), p0-pMN in *pcdh19* morphant embryos (N=3, n=10), p0-p0 in *cdh2* morphant (N=2, n=15), p0-pMN in *cdh2* morphant (N=3, n=12). (E) pMN domain at the neural tube stage, visualized by *TgBAC(olig2:GFP)*, in control or *cdh11* morphant embryos. The cross-sections in the *cdh11* morphants show the dorsal mislocalization of a pMN cell (white arrow) or a *olig2:GFP* negative cell mixed in the pMN domain (yellow arrow). Scale bars are 20 µm. (F) Quantification of mixed cells in the pMN domain in embryos injected with standard control morpholino (N=3, n=15) or *cdh11* MO (N=3, n=21). For (F,H), the boxes within the box plot represent 25^th^, 50^th^, and 75^th^ percentile of the data. *** represent p-value < 0.001 (t-test). (G) p0 domain at the neural tube stage, visualized by *TgBAC(dbx1b:GFP)* in wildtype, *cdh2* mutant, or embryos injected with *cdh2* morpholino in 1 cell of 8-cell stage embryos. Scalebars are 30 µm. (H) Quantification of mixed cells in p0 domains in *cdh2* homozygote mutants (N=2, n=6) and their wildtype or heterozygote siblings (N=2, n=15), embryos injected in 1 cell at 8-cell stage with standard control morpholino (N=3, n=9), or *cdh2* morpholino (N=3, n=17). (I) A cartoon diagram summarizing the working model for homotypic preference between the pMN and p0 cells. The pMN cells utilize Cdh11 while the p0 cells utilize Cdh2 and the Cdh2-Pcdh19 complex to achieve homotypic preference.

To analyze the homotypic preference of p0 cells against pMN cells, we examined the roles of Pcdh19 and Cdh2, the two adhesion molecules enriched in p0 cells relative to pMN cells. As was the case for *pcdh19* knockdown in pMN-p3 heterotypic adhesion, loss of Pdh19 also leads to a Cdh2-dependent increase of pMN-p0 heterotypic adhesion (Figure 6D). However, while Pcdh19-deficient p3 cells showed no homotypic preference, Pcdh19-deficient p0 cells did not entirely lose their preference for binding to each other (Figure 6C). We hypothesized that this is due to higher levels of Cdh2 in the p0 cells than the pMN cells (Figure 4C,N). Even without the distinct adhesion mode caused by Pcdh19-Cdh2 complex formation, higher levels of Cdh2 in p0 cells could still create a preference for p0-p0 homotypic contact over the pMN-p0 contact. Consistent with this, loss of Cdh2 abolished the homotypic preference of p0 cells against pMN cells (Figure 6C). Without Cdh2, both p0-p0 homotypic adhesion and p0-pMN heterotypic adhesion were negligible (Figure 6D). We conclude that the homotypic preference of p0 cells against pMN cells is achieved primarily by higher levels of Cdh2-based adhesion, with a minor contribution by Pcdh19.

To verify the roles of Cdh11 and Cdh2 in patterning of the pMN and p0 domains *in vivo*, we used morpholinos or previously reported mutants to complement our CRISPR screen. Consistent with the phenotype of *cdh11* CRISPR mutants, *cdh11* morphant embryos also exhibit disrupted pMN domains (Figure 6E,F). The patterning phenotypes include mixing of *olig2:GFP* negative cells within the *olig2:GFP* positive domain, or dorsal mislocalization of pMN cells (Figure 6E). Since pMN and p0 domains are not adjacent to each other at the neural tube stage, the most important pMN-p0 interactions *in vivo* take place during the neural keel stage, when the converging p0 cells appear to squeeze the pMN cells at the dorsal tip of the wedge-shaped pMN domain (Figure 1K). At this stage, lack of homotypic preference of pMN cells versus p0 cells would result in the detachment of pMN cells from the rest of the pMN domain and mislocalization of pMN cells dorsally. Indeed, in the *cdh11* morphant or CRISPR mutant embryos we frequently observed dorsal mislocalization of pMN cells caused by detachment of pMN cells from the pMN domain at the neural keel stage (Figure 6E and Movie S6).

To verify the p0 patterning phenotype in our *cdh2* CRISPR mutant, we used either *cdh2* morpholinos or a classic *cdh2* mutant, parachute (Jiang et al., 1996). In the *cdh2* mutant, the p0 pattern also exhibited severe defects (Figure 6G,H). However, the interpretation of the patterning phenotype in the *cdh2* mutant is complicated by the accompanied defect in morphogenesis. Due to the delay in convergent movement, the *cdh2* mutant embryo is known to have a “mushroom” shaped spinal cord (Lele et al., 2002) (Figure 6G). The morphogenesis defect is phenocopied by our *cdh2* CRISPR mutant and *cdh2* morphant. The p0 domain is particularly affected as it is positioned at the region connecting the “stalk” and the “hat” of the mushroom shaped spinal cord (Figure 6G). To separate patterning and morphogenesis defects, we performed a milder perturbation by injecting *cdh2* morpholino in 1 cell of the 8-cell stage embryo. We only analyzed embryos whose overall shape of the spinal cord were not affected by this mild perturbation, and still observed significant disruption of the p0 pattern (Figure 6G,H). Together, our results suggest that the differential expression of *cdh11* and *cdh2* mediate the homotypic preference of pMN and p0 cells *ex vivo*, and also the cohesion of pMN and p0 domains *in vivo* (Figure 6I).

### Shh co-regulates cell fate and adhesion code in the ventral spinal cord

To generate a cell-type-specific adhesion code, the expression of adhesion molecules and cell fate specification need to be closely coupled. We speculated that the adhesion molecules must share some common upstream regulators with the cell fate specification process. Shh is a key morphogen secreted from the ventral side of the spinal cord, and instructs specification of ventral neural progenitor fates (e.g., p3 and pMN cells) in a dose-dependent fashion (Briscoe and Ericson, 2001; Park et al., 2004). We therefore set out to perturb Shh signaling and observe the expression patterns of both the cell fate markers for pMN and p3 cells (i.e., *olig2* and *nkx2.2a*) and their corresponding adhesion molecules (i.e., *cdh11* and *pcdh19*).

To down-regulate Shh signaling, we treated embryos with cyclopamine, a chemical inhibitor of Smoothened, a downstream activator of Shh (Chen et al., 2002). We began treatment of 100 µM cyclopamine at 4 hours post fertilization, prior to the expression of endogenous Shh. To up-regulate Shh signaling, we globally over-expressed it by injection of 90 pg of *shha* mRNA into one-cell-stage embryos. In wildtype embryos, *olig2* and *cdh11* expression are highly correlated (Figure 4F,K,N). Under cyclopamine treatment, the expression level of *olig2* and *cdh11* are both significantly reduced (Figures 7A-D, S4A). Conversely, when Shh is hyperactivated, both *olig2* and *cdh11* significantly expand their expression domains (Figure 7E-H, S4B). Interestingly, even under Shh down- and up-regulation, although the positions and levels of *olig2* and *cdh11* expression were both shifted, they still shared the same relative pattern with overlapping peak positions and a slightly broader domain for *cdh11* (Figure 7A-H). Furthermore, expression of *olig2* and *cdh11* remained correlated at the single cell level under both perturbations (Figure S4 C-F). This suggests that *olig2* and *cdh11* respond to Shh signaling in a similar but not identical fashion.

**Figure 7.**
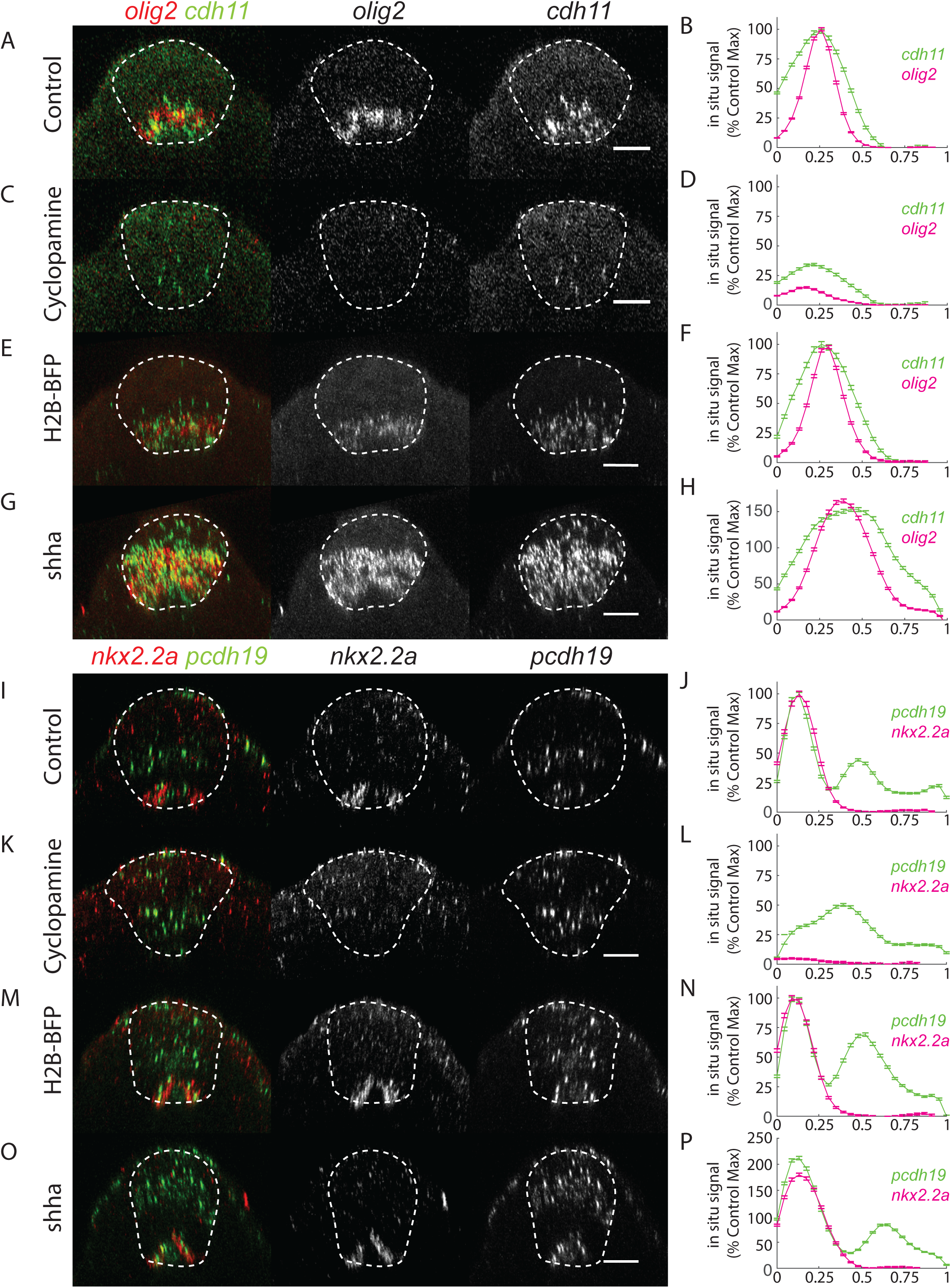
Shh co-regulates cell fate and adhesion code in the ventral spinal cord. (A, C) Cross-section of a 11-somite stage embryo treated with vehicle control (A) or 100 μM cyclopamine (C), stained with *in situ* HCR probes against *olig2* and *cdh11*. All scale bars in this figure are 20 µm. (B, D) The ventral-to-dorsal *in situ* HCR profile of *olig2* and *cdh11* from embryos treated with (B) vehicle control (16 Embryos, 1596 sections quantified) or (D) 100 μM cyclopamine (14 Embryos, 1414 sections quantified). (E, G) Cross-section of a 11-somite stage embryo injected with 90pg of H2B-EBFP2 mRNA (E) or 90 pg of *shha* mRNA (G), stained with *in situ* HCR probes against *olig2* and *cdh11*. Images are the maximal intensity projection over 1.79 um wide slab. (F, H) The ventral-to-dorsal *in situ* HCR profile of *olig2* and *cdh11* from embryos injected with (F) 90pg of H2B-BFP mRNA (16 Embryos, 1616 sections quantified) or (H) 90pg of Shha mRNA (15 Embryos, 1515 sections quantified). (I, K) Cross-section of a 11-somite stage embryo treated with vehicle control (I) or 100 μM cyclopamine (K), stained with *in situ* HCR probes against *nkx2.2a* and *pcdh19*. (J, L) The ventral-to-dorsal *in situ* HCR profile of *nkx2.2a* and *pcdh19* from embryos treated with (J) vehicle control (10 Embryos, 1010 sections quantified) or (K) 100 μM cyclopamine (11 Embryos, 1111 sections quantified). (M, O) Cross-section of a 11-somite stage embryo injected with 90pg of H2B-EBFP2 mRNA (E) or 90 pg of *shha* mRNA (G), stained with *in situ* HCR probes against *nkx2.2a* and *pcdh19*. (N, P) The ventral-to-dorsal *in situ* HCR profile of *nkx2.2a* and *pcdh19* from embryos injected with (N) 90pg of H2B-BFP mRNA (11 Embryos, 1111 sections quantified) or (P) 90pg of *shha* mRNA (13 Embryos, 1313 sections quantified).

Next, we examined the expression pattern of the p3 cell marker *nkx2.2a*, and its corresponding adhesion molecule, *pcdh19*, under Shh perturbations. In wildtype embryos, *pcdh19* is expressed in two stripes, with the ventral stripe overlapping with the *nkx2.2a* expressing domain (Figures 4H,J,7I,J). Cyclopamine treatment completely abolished *nkx2.2a* expression. Interestingly, the *pcdh19* expression switched from two stripes to one broader stripe centered near the original dorsal stripe, likely due to the disappearance of the ventral stripe (Figure 7K,L). When Shh is hyperactivated, both *nkx2.2a* and the ventral stripe of *pcdh19* significantly increased the transcript level by 1.5-2 fold, and the dorsal stripe of *pcdh19* expanded more dorsally than in control embryos (Figure 7M-P). Again, the relative patterns of *nkx2.2a* and *pcdh19* are preserved through up- and down-regulation of Shh, suggesting shared regulatory logic between *nkx2.2a* expression and the ventral expression of *pcdh19* by Shh. These findings bridge the morphogen gradient-based French Flag Model with the differential adhesion-based cell-sorting mechanism, and suggest a close interplay between these two classic models during neural tube construction.

## DISCUSSION

How spatial patterns form reproducibly in a morphogenetic tissue is an important open question. By studying the patterning of three neural progenitor domains in the zebrafish spinal cord, we identified a cell-type-specific adhesion code in three cell types that allows them to form stronger homotypic adhesion and domain cohesion during morphogenesis. Following the exposure to morphogen signal gradients (e.g., Shh), neural progenitors not only begin committing to different fates by expressing distinct cell fate regulators, but also express adhesion molecules differentially to mediate cell sorting. We conclude that the interplay between mophogen-based pattern formation and differential adhesion-based sorting allows robust pattern formation in the zebrafish spinal cord despite imprecise morphogen signaling and major cellular movements during tissue morphogenesis.

### Completeness of the adhesion code

Our work reveals an adhesion code composed of *cdh2*, *cdh11*, and *pcdh19* that mediates patterning of three neural progenitor domains in the zebrafish spinal cord. This finding is likely a subset of a more complex patterning system involving additional adhesion molecules. For example, p0 cells exhibit homotypic preference over pMN cells due to higher levels of Cdh2. Given that all dorsal cell types express a high level of Cdh2, this is likely a more general property for all dorsal cell types to allow them to adhere to each other against ventral cell types. Within the dorsal cell types, the differences of Cdh2 would not be sufficient to mediate sorting of neighboring domains. Yet the need to maintain domain cohesion is perhaps even stronger in the dorsal half of the spinal cord than in the ventral half, because of the larger range of morphogenetic movements it undergoes. In addition, *cdh11* expression is broader than the pMN domain. Therefore, *cdh11* is likely also expressed in domains next to the pMN domain, such as the p2 domain. While *cdh11* mediates homotypic preference of pMN cells against dorsal cell types like p0 cells, it may not be sufficient to mediate the homotypic preference of pMN cells against p2 cells. It is likely that additional adhesion molecules contribute to homotypic preference and sorting within dorsal neural progenitors or between p3, pMN, p0 cells and other neural progenitor cell types.

### Coordination between cell fate specification and the adhesion code

The adhesion code is closely coupled to cell fate specification. At the ventral neural tube, specification of ventral neural progenitors (e.g., pMN and p3 cells) and the expression of their corresponding adhesion molecules (e.g., *cdh11* and *pcdh19*) are both instructed by Shh, but the detailed mechanisms are still unknown. One simple model to explain this coordination would be that the cell fate regulator directly mediates the expression (or lack of expression) of specific adhesion molecules. This would require that the expression of the cell fate regulators precedes that of the adhesion molecules. Alternatively, the expression of the cell fate regulator and the adhesion molecules could both be induced by a common upstream transcription factor, or might each respond independently to morphogen signaling. In this case, the adhesion molecule might be expressed earlier than the cell fate regulator. We noticed that *pcdh19* expression precedes the expression of *nkx2.2a*, the p3 cell fate marker, suggesting that the coupling between p3 fate specification and *pcdh19* expression may not be as simple as a direct response of *pcdh19* to *nkx2.2a*. We also note that adhesion-based sorting could influence a cell’s exposure to morphogen and thus indirectly affect cell fate specification. The gene regulatory network underlying the expression pattern of adhesion molecules remains to be fully characterized.

### The adhesion code enables homotypic preference simultaneously in three cell types

The mechanics of cell sorting have attracted much attention since Steinberg proposed the differential adhesion hypothesis 50 years ago (Steinberg, 1970). Valuable biophysical insights have come from studies of cadherin-based cell sorting using cell types expressing different levels of a single cadherin (e.g., E-cadherin) (Foty and Steinberg, 2005; Krieg et al., 2008; Maître et al., 2012). Interestingly, a recent study suggested cell sorting is more robust under high heterotypic interfacial tension (Canty et al., 2017). This condition requires the interfacial tension between all homotypic contacts to be lower than the heterotypic contacts, a phenomenon similar to the homotypic preference described in our work. When cells rely only on different levels of a single cadherin to sort, the cell type with lowest level of cadherin would fail to exhibit homotypic preference, and the system would fail to sort reliably. Here we performed a detailed biophysical characterization to address how three types of cells can simultaneously achieve homotypic preference through differential expression of three adhesion molecules. We found that multiple adhesion specificities can be created through combinations of type I and type II cadherins (e.g. Cdh2 vs Cdh11) as well as through adhesive-specificity switching by protocadherin (e.g Cdh2 vs. Cdh2-Pcdh19).

### Interplay between the French Flag and the Differential Adhesion models

The French Flag model and the Differential Adhesion model are two classic proposals for how tissue-scale patterns emerge in development. While both models provide valuable insights in many biological contexts, each model has its own limitations. When a morphogen gradient instructs tissue patterns, errors from biochemical fluctuations are inevitable, and cell neighbor exchanges during morphogenesis make it difficult to explain the emergence of robust patterns. On the other hand, while the Differential Adhesion model allows distinct cell types to self-organize and sort into spatially separated domains, by itself it is unable to generate an oriented and positioned pattern. Differential adhesion based sorting is also sensitive to the initial conditions: if cells of the same type are specified far apart, it may take a long time to complete cell sorting, and cells may be trapped in a local energy minimum when they are completely surrounded by different types of cells. If morphogen gradients control differential adhesion, this may combine the advantages of both models. In this scenario, a morphogen gradient specifies an initial pattern of cells with distinct adhesion properties, such that cells of the same adhesion properties are near to each other. The cells then self-organize into well separated domains, overcoming any imperfections in the initial pattern laid down by the morphogen, and remaining organized in domains despite the neighbor exchange that occurs during tissue morphogenesis. Our data suggests that such a combined mechanism allows robust neural progenitor patterning in the zebrafish spinal cord, and that similar mechanisms may offer a general solution for precise pattern formation in morphogenetic tissues.

## ACKNOWLEDGMENTS

We thank the members of the Megason and Heisenberg labs for critical discussions of the work. Rebecca Ward, Fengzhu Xiong, Anna Kicheva, Akankshi Munjal, Natasha O’Brown have given valuable comments on the manuscript. We thank Fengzhu Xiong for training on *in toto* imaging and the initial characterization of the sorting process in the zebrafish spinal cord, Ian Swinburne for help with design of CRISPR screening strategies and general technical discussions, Daniel Wagner for assistance in analysis of RNA-seq data, Jana Slovakova and the staff at the Bioimaging facility of IST Austria for help with micropipette aspiration assays, Allon Klein for insights in single cell dissociation protocols, Kana Ishimatsu and Natasha O’Brown for help with *in situ* hybridization, Sandy Nandagopal for advice about multiplex *in situ* HCR, Sharon Cooper for insights in protocadherin 19 regulation, Dante D’India, Andrew Murphy, Verena Meyer and Deborah Godard for fish care. Although not included in this manuscript, Danylo Lavrentovich, Preeti Sahu, Lisa Manning, and Jennifer Schwartz have implemented computational simulations that were helpful for our insights in cell sorting. We thank the laboratories of Bruce Appel, Scott Holley, James Jontes, Darren Gilmour for sharing the transgenic zebrafish strains, and the laboratory of Chenghua Gu for sharing their cryostat for frozen sectioning. T.Y.-C.T. was supported by the Damon Runyon Cancer Foundation and the NICHD K99 pathway to independence fellowship (1K99HD092623). The collaboration between the Megason and the Heisenberg labs was generously supported by the travelling fellowship of the Company of Biologists, and the Collaborative Research Grant from the Burroughs Wellcome Foundation to T.Y.-C.T.. T.Y.-C.T. and S.G.M. were supported by NIH Grant R01GM107733. T. C.-C. and H.K. were supported by NIH Grant R01NS102322. C.-P. H. was supported by an ERC advanced grant (MECSPEC)

## MATERIALS AND METHODS

### Zebrafish Strains

Zebrafish husbandry and experimental usage were approved by the Harvard Medical School Animal Care and Use Committee (Protocol number: 04487) and performed in accordance with the guidelines. Zebrafish (*Danio rerio*) AB and TL strain were used in this manuscript. Adult zebrafish were maintained on a 14 h light / 10 h dark cycle at ∼28°C. Zebrafish embryos were obtained by crossing male and female adults aged 3-24 months. Embryos are collected 15 min to 1h after egg laying and raised at various temperature between 23 to 28.5°C (as described in corresponding assays). Gender at the developmental stages studied is not yet determined and therefore cannot be distinguished.

The *cdh2*^tm101^ allele contains a nonsense mutation leading to a premature stop codon in the extracellular domain of *cdh2* (Lele et al., 2002). The *pcdh19*^os51^ allele contains a 10 bp deletion, resulting in a premature stop codon (Cooper et al., 2015). *Cdh2* and *pcdh19* homozygous embryos were generated by inbreeding heterozygous adults. *cdh2* mutants were identified by morphology of the spinal cord at 10 somite stage and gross morphology of the whole embryo after 2dpf. Mutant sequence is further confirmed by PCR amplification followed by sequencing of the amplicon. Pcdh19 mutants were identified by differences in size of the amplicon through PCR amplification (166 bp for wildtype and 156 bp for mutant. The difference was detectable on a 3% agarose gel (Cooper et al., 2015). Primers used for genotyping are as follows; *cdh2*: forward 5’agcttccacctgctgttca3’, reverse 5’ataaaacctttaaaataatttgatcgc 3’, *pcdh19*: forward 5’cgcgtgaagacagacatcaacagg3’, reverse 5’cgttacgttggctatctttgtgcc3’.

The following transgenic zebrafish were used. *TgBAC(olig2:GFP)* (Shin et al., 2003), *TgBAC(nkx2.2a:memGFP)*(Kirby et al., 2006; Ng et al., 2005), *TgBAC(dbx1b:GFP)* (Kinkhabwala et al., 2011), *TgBAC(cdh2:cdh2-Cherry)* (Colak-Champollion et al., 2019), *Tg(actb2:memCherry2)^hm29^* (Xiong et al., 2013), *Tg(actb2:memCitrine-Citrine) ^hm30^* (Xiong et al., 2014). Transgenic embryos were identified by fluorescence.

### Morpholinos and CRISPR

For experiments using morpholino against *cdh2* and *cdh11*, embryos are injected with either 2.3nl of 0.25 mM morpholino, or 1.1nl of 0.5mM morpholino (corresponds to 2.1 ng). For experiments using morpholino against pcdh19, embryos are injected with either 2.3 nl of 0.5mM morpholino, or 1.1 nl of 1mM morpholino (corresponds to 4.2 ng). We used sequences of morpholino against *cdh2*, *cdh11*, and *pcdh19* that were previously validated for efficacy (Clendenon et al., 2009; Emond et al., 2009; Lele et al., 2002). The sequences are *cdh2* MO (TCTGTATAAAGAAACCGATAGAGTT), *cdh11* MO (CCCCATCAGGTAGAGTCTGCT TCCT), *pcdh19* MO (TCCTTGGAATGCATTGTACCTGTTGA). The *cdh2* morphant phenocopied the *cdh2* mutant (Lele et al., 2002). The *cdh11* and *pcdh19* MO were validated by immunostaining and immunoblotting (Clendenon et al., 2009; Emond et al., 2009).

To generate CRISPR mutant zebrafish embryos, we followed the previously described protocol (Gagnon et al., 2014). In brief, 3-4 guide RNAs for each target were designed using the online tools ChopChop (https://chopchop.cbu.uib.no/), synthesized as DNA, and in vitro transcribed. Sequences of guide RNAs can be found in the Extended Experimental Procedures. Guide RNAs targeting the same gene were pooled and combined with purified Cas9 protein to inject into 1 cell stage embryos. A guide RNA-Cas9 injection cocktail contains 50∼100ng/μl of the pooled guide RNAs and 5.6 μM of Cas9 protein, with an injection volume of 2.3nl. Injected embryos were kept at 28°C until shield stage (6hpf) and then moved to 23°C for imaging at 10 somite stage the next day. The lack of pigment in the tyrosinase CRISPR mutant was used to assess the efficacy of the Cas9 protein and also served as a negative control for neural progenitor patterning.

### Confocal imaging

For time-lapse microscopy to observe the formation of neural progenitor domains *in vivo*, we adapted the previously described protocols for *in toto* imaging (Megason, 2009; Xiong et al., 2013). Specifically, tailbud stage embryos were dechorionated, and mounted in a dorsal mount with the dorsal side closest to the objective (Megason, 2009). Confocal z-stacks were taken on a Zeiss LSM710 laser scanning confocal microscope, using a C-Apochromat 40X 1.2 NA objective or a Plan-Apochromat 20X 1.0 NA objective. In a typical experiment, we imaged a 265 µm * 265 µm square region, with a depth of 120 µm. This will cover about two-thirds of the neural plate along the medial-lateral axis, and is sufficient to include the future p0 cells. This volume also covers the whole neural keel and neural tube as the neural plate thickens dorsally. Typically, this volume will span the 2^nd^ to 6^th^ somite and the entire neural tube within this region. We kept the embryos at 22°C to enable better temporal resolution for cell tracking. Under this temperature, a frame rate of 5-8 minutes was sufficient for single cell tracking.

To evaluate neural progenitor patterning at the neural tube stage, 8-11 somite stage embryos were mounted on the dorsal mount, with the dorsal side of the neural tube closest to the objective. Confocal z-stacks were taken in a 265 * 265 * 120 µm volume that contains the 2^nd^ to 6^th^ somite and the entire neural tube within this region. Fixed embryos stained with *in situ* HCR probes were mounted in a similar fashion.

### Whole mount *in situ* hybridization

Synthesis of RNA probe and in situ hybridization were performed as previously described (Thisse and Thisse, 2008). Probes against *cdh11* and *pcdh19* were prepared with a DIG RNA labeling mix (Roche, Cat. No. 11277073910) using primers cdh11: forward 5’GAATTTAATACGACTCACTATAGGGAGACC ttaagagttgtcatcgacggagtctt3’, reverse 5’gtcaaagatgtgaatgataacgctcc3’, *pcdh19:* forward 5’ GAATTTAATACGACTCACTATAGGGAGACC aagcacgatgtccttcagtc 3’, reverse 5’gctgtcaagtgcaaaaggg3’, and detected with anti-DIG-AP antibody (1:5000; Sigma-Aldrich, Cat. No. 11093274910) and NBT/BCIP stain (1:1000; Roche, Cat. No. 11383213001 and 11383221001). Stained embryos were imaged with a 1x 0.25 NA objective on a Olympus MVX10 microscope equipped with a Axiocam MRc color camera (Zeiss).

### Multiplex *In situ* hybridization chain reaction (HCR)

We followed the protocols suggested by Molecular Instruments. Embryos were fixed with 4% paraformaldehyde in phosphate buffered saline solution (PBS) at 4°C overnight, washed with PBS, and dehydrated with serial methanol washes (25%-50%-75%-100% methanol in PBS) before storage in −20°C freezer. On the first day of staining, embryos were rehydrated by serial dilution of methanol (75%-50%-25% of methanol in PBS), washed with PBS, and pre-hybridized in hybridization buffer (Molecular Instrument). Embryos were then incubated in 200 μl of hybridization solution containing 1pg of probes overnight at 37°C. The next day, embryos were washed 4 times in wash buffer (Molecular Instruments) followed by two washes in 5x SSCT (5x of SSC buffer and 0.1% of tween 20). For the pre-amplification step, embryos were incubated in amplification buffer (Molecular Instruments) for more than 30 minutes. At the same time, hairpin mixtures were prepared by heating 12 pmol of hairpin 1 and 2 for each sample to 95 °C for 90 seconds, and snap-cooled in the dark. The snap-cooled hairpins were added to 200 μl of amplification buffer. Embryos were incubated in the hairpin mixture at room temperature overnight in the dark. On the third day, embryos were washed more than 4 times in 5xSSCT and either stored in 4°C or mounted for confocal microscopy.

### Immunostaining and in situ hybridization of frozen-sectioned embryos

Embryos were fixed with 4% paraformaldehyde in PBS at 4°C overnight. After PBS washes, embryos were incubated with 30% sucrose in PBS at 4°C overnight for cryoprotection. Cryoprotected embryos were embedded in O.C.T. embedding media (Tissue-Tek No. 4583) and frozen by dry ice. Embedded samples were sectioned at a thickness of 14 μm and placed on superfrost slides (VWR) for following immunostaining or *in situ* HCR.

For immunostaining, embryo sections were rinsed with PBS and permeabilized with PBSTT (PBS, 0.5% Triton-X and 0.1% Tween20) at room temperature for more than 1 hour. Permeabilized embryo sections are blocked with blocking solution (PBS, 0.1% Triton-X, 0.1% Tween20, 1% DMSO, 10% Goat serum) for more than 2 hours at room temperature and incubated with primary antibody solution (1:200 of Cdh2 antibody in blocking solution) overnight at 4°C. On the next day, slides were washed with PBSTT at room temperature for at least 4 times then incubated with secondary antibody (1:400 of secondary antibody in blocking solution) at 4°C overnight.

For *in situ* hybridization with HCR, protocols were similar to whole-mount embryos, with the exception that embryo sections were initially dehydrated by serial ethanol washes (50%-70%-100%) before pre-hybridization as previously described (O’Brown et al., 2019). For both immunostaining and *in situ* HCR samples, slides were mounted in VECTASHIELD antifade mounting media (Vector laboratories H-1000) after staining and sealed with nail polish.

### Image analysis

#### Analysis of dorsal-ventral intensity profile of fluorescent reporters and fluorescent in situ hybridization

Custom-made scripts in Matlab (Mathworks) were used to allow semi-automatic analysis of the ventral-to-dorsal intensity profile of fluorescent reporters and fluorescent *in situ* hybridization. Briefly, each of the confocal z-stacks spans 265 µm (800 pixels with 0.332 µm/pixel resolution) along the anterior-to-posterior axis, corresponding to the length of 5 somites. We chose 15 cross sections that are 16.6 µm (50 pixels) apart along the anterior-posterior axis, and manually defined 4 points in each section. These first 2 points established the ventral-to-dorsal axis, corresponding to the ventral-most point of the spinal cord (i.e., the ventral surface of the medial floor plate cell) and the dorsal-most point of the spinal cord (i.e., the dorsal surface of the roof plate cells). The next 2 points specified the medial-lateral width of the domain to analyze. We chose the left-most and right-most point of the p3 domain (Figure S1A). This width of the p3 domain is typically ∼20 µm. In the case when the p3 domains were not imaged, we estimated a width of about 20 µm. Based on the manually defined ventral-to-dorsal axis and the width, 24 boxes were drawn (Figure S2B) and the average fluorescent intensity within each box calculated. The 24 intensity measurements along the ventral-to-dorsal axis were then plotted as a single ventral-to-dorsal intensity profile (Figure S1C). For the cross-sections between these manually analyzed sections, the ventral-most points and dorsal most points of each section were defined by interpolation from the manually labelled positions. The ventral-to-dorsal intensity profiles were calculated from every cross section that are 1.6 µm apart. We typically excluded the anterior 15 µm and posterior 15 µm regions of the confocal z-stack, resulting in ∼140 cross sections analyzed from each embryo. The final ventral-to-dorsal intensity profiles (e.g., Figure 1D) were the average intensity profile over all profiles from all embryos measured. For comparison of the ventral-to-dorsal intensity profiles between perturbed embryos and control embryos (e.g., Figure 7), we only included results when perturbed embryos and control embryos were stained and imaged in the same experiment, and the intensity profiles were normalized to the maximal intensity of the control embryos.

#### Quantification of mixed cells during neural progenitor patterning

For each confocal z-stack we acquired, the field of view spanned an A-P distance of 265 µm, corresponding to the position of the 2^nd^ to 6^th^ somites at the neural tube stage. We counted the number of mixed cells (defined below) in 31 equal distance sections. Each section is 8.86 um apart, roughly equals to 1 cell diameter. Within each cross-section, a mixed cell typically falls within one of the three categories. (1) a GFP-negative cell is mixed within the GFP positive domain and appear as a hole in the domain (e.g., Figure S1H and 6E). In this case, one can draw a line from the ventral most point of the neural tube, passing the GFP-negative cell, and this line will cross the GFP-positive cell twice (once ventral to the GFP-negative cell and once dorsal to the GFP-negative cell). (2) a GFP-negative cell breaks the GFP positive domain into 2 disconnected domains. One can connect the two GFP-positive domains by replacing this one GFP-negative cell to a GFP-positive cell (e.g., Figures 1E,I, S1I,J, and 6G). (3) A GFP-positive cell is mislocalized away from the GFP-positive domain (Figure 6E). By definition, this is a special case of (2), because there is also a GFP-negative cell that separates the GFP positive domain and the one mislocalized GFP-positive cell into 2 disconnected parts. The distinction is that in (2), all the GFP-positive cells are at the proper position, but the GFP-negative cell intrudes the GFP-positive domain and breaks it into 2 parts. And in (3), there is no intrusion of GFP-negative cells but instead the GFP-positive cell intrudes the neighboring GFP-negative domain. In this scenario, we counted how many GFP-negative cells are between the GFP-positive domain and the one mislocalized GFP-positive cell as the number of mixed cells. We believe this definition is meaningful. For example, a GFP-positive cell that is mislocalized to a distance of three cells away is a more severe phenotype than a GFP-positive cell that is mislocalized to a distance of two cells away.

We applied a different criterion to count the number of mixed cells along the p3-pMN boundary. This is because the p3 cells are more sparsely distributed under certain perturbations, and there may be only one mislocalized p3 cell in some cross sections. In the case of p3-pMN boundary, we examined the relative position of p3 and pMN cells by simultaneous imaging of the p3 and pMN domains in the transgenic reporter embryos TgBAC(*nkx2.2a:memGFP);TgBAC(olig2:dsRed)*. In the wildtype embryos, all the p3 cells should locate at a more ventral position than pMN cells and are physically closer to the Shh-producing medial floor plate cells. Therefore, we counted as one mixed cell for the p3-pMN boundary for each p3 cell that is positioned more dorsally from the medial floor plate than another pMN cell. Under this definition, if there is only one p3 cell in the cross-section, but it is positioned more dorsally than another pMN cell, we still count this as a mixed cell.

#### Single cell intensity analysis of multiplex *in situ* HCR

Embryos of transgenic zebrafish *Tg(actb2:memCitrine)* were used to visualize cell membranes. For each embryo, two HCR probes were used and visualized with amplifiers conjugated to either AF546 or AF633. In each confocal z-stack, we chose a volume covering 265 µm along the anterior-to-posterior axis, corresponding to the 2^nd^ to 6^th^ somite, and imaged the whole depth of the spinal cord within this region. To pick single cells, we selected 31 equal-distance sections along the A-P axis. Each section is 8.85 µm apart. We placed a seed at the center of each cell using the membrane localized Citrine signal as a guide, and quantified the average fluorescent intensity of AF546 and AF633 in a 3.5 * 3.5 * 3.5 (µm^3^) region centered at the manually defined spots (Figure S3E). We only chose cells with sufficient margin around the center for quantification to avoid averaging signals from two cells within the same volume. For comparison between perturbed embryos and control embryos (Figure S4), we only included results when perturbed embryos and control embryos were stained and imaged in the same experiment, and the intensity profiles were normalized to the maximal intensity of the control embryos.

### Micropipette aspiration assays

#### Dual pipette aspiration assay

We followed the detailed protocol previously described with the following modifications (Biro and Maître, 2015). As the neural progenitor cells were smaller than the cells used in previous experiments, the diameter of the micropipettes was adjusted to 4 ± 0.5 μm. After coating of micropipette with heat-inactivated serum, we soaked the micropipettes in PBS for over an hour to ensure air was purged from the tip of the micropipette by capillary effect. For each experiment, 10-15 embryos were manually dechorionated and dissected. We made the wire loop for dissection by inserting a loop of thin stainless-steel wire through a glass capillary tube mounted on a halved chopstick. For each embryo, we dissected out the tailbud region and the embryonic structures anterior to the hindbrain, leaving the trunk region of the embryo. The trunk segments of all dissected embryos were dissociated through manual pipetting in calcium free PBS, and plated on a glass-bottom dish containing 0.9X of DMEM/F12 media. We typically started the pipette aspiration measurement within 20 minutes of cell dissociation. For each pipette aspiration experiment, two fluorescent positive cells (of the same or different types) were chosen. The membrane GFP signal of the p3 cells from the *TgBAC(nkx2.2a:memGFP)* reporter can be clearly distinguished from the cytoplasmic GFP signal of the pMN or p0 cells from the *TgBAC(olig2:GFP)* or *TgBAC(dbx1b:GFP)* reporters (Movie S5). To distinguish the pMN and p0 cells, we injected 1ng of Texas Red conjugated 3kD dextran (Thermo Fisher No. D3329) into one cell stage embryo of either the *TgBAC(olig2:GFP)* fish or *TgBAC(dbx1b:GFP)* fish (Movie S6). This allowed us to select the *olig2:GFP* or *dbx1b:GFP* positive cells through the presence of fluorescent dextran. Although we did not see a bias in the triplet assay between the dextran-injected and non-injected cells, we alternated the injection between *TgBAC(olig2:GFP)* or *TgBAC(dbx1b:GFP)* between experiments. Once the two selected cells formed a doublet, we kept track of the contact formation time. Our pilot experiments showed that the average adhesion force measured after 3 minutes of contact formation was stronger than 1 minute of contact formation, but not much weaker than 10 minutes of contact formation (Figure S2A). Therefore, we allowed the cell doublets to form contact for 3 minutes before initiating the pipette pulling sequence. After 3 minutes, we started preparing the pulling assay by increasing the holding pressure of the left pipette to 80 Pascal, grabbing the right cell of the doublet with the right pipette, and switch the pipette motor to a computer-controlled mode. For each pulling measurement, we increased the pressure sequentially, and pulled by moving the right pipette to the right by 20 μm at a speed of 20 μm/sec (Movie S5). A typical sequence of aspiration pressure is as follow (unit: Pascal): 2→5→8→12→16→20→30→40→60→80→100→130→160→200→240. With a pipette diameter of 4 μm, this will correspond to a pulling force (nN) of 0.25→0.62→1.0→1.5→2.0→2.5→3.8→5.0→7.5→10.0→12.6→ 16.3→20.1→25.1→30.2

To avoid pulling the same doublet too many times, we started the pulling sequence at a force closer to the average of the doublet estimated from pilot experiments. For example, for pMN-pMN doublets which exhibit an average adhesion force of 7nN (Figure 2B), we skipped steps in the smaller force regime and a typical sequence will instead be 10→20→30→40→60→80→100→130→160→200→240. In a pipette with 4 μm diameter, this corresponds to a pulling force (nN) of 2.5→3.8→5.0→7.5→ 10.0→12.6→16.3→20.1→25.1→30.2. We recorded the average pressure between the last pressure that the doublet remains adhered and the first pressure that the doublet breaks. For each doublet we measured, we also estimated the contact area (A) based on the length of the contact (L) using equation, Area = π*(L/2)^2^. As we did not observe a significant difference of contact size among the 6 combinations of doublets (Figure S2B), we did not show data normalized by the contact area in the main figure (Figure 2B).

#### Triplet assay

The micropipettes were prepared with identical protocol as the dual pipette aspiration assay. Here we use the pMN-p0-p0 triplet as an example (triplet configuration corresponds to Movie S6). Two p0 cells and one pMN cell were chosen based on fluorescence. All three cells started as isolated cells. At the beginning of the assay, left pipette held on to one pMN cell while right pipette held on to one p0 cell. The remaining p0 cell was moved by stage motor to the center of the field of view. The pMN cell held by the left pipette and the p0 cell held by the right pipette approached the p0 cell at the middle simultaneously. When both contacts were made, we left the triplet unperturbed for at least 3 minutes to allow contact formation. After that, we started to prepare for pipette aspiration. We increased the aspiration pressure in both pipettes to an aspiration force beyond 10nN to avoid detachment of cells from either pipette during pulling. The right pipette was set to a computer-controlled mode with a speed of 20 μm/sec. And the pipette pulling initiates after at least 3 minutes of initial contact. Care was taken to alternate the left-right order of the 2 contact types (e.g., p0-p0-pMN followed by pMN-p0-p0). While we always pulled on the cell at the right of the triplet, we did not see a significant bias that the right cell was more or less preferred in the outcome of the assay (Figure S2C). We’ve observed examples where one of the two contacts was lost before pulling, and the cell in the middle automatically chose to adhere to only one cell. In this case, if the contact was lost within 1 minute of the initial contact, we concluded the initial contact formation failed and we would reset the experiment. If the contact was lost after 1 minute of the initial contact, we scored the remaining contact as the preferred one.

#### FACS sorting of neural progenitors and RNA-seq

To obtain the transcriptomes of p3, pMN, or p0 cells, we made use of the *TgBAC(nkx2.2a:memGFP)*, *TgBAC(olig2:GFP)*, or *TgBAC(dbx1b:GFP)* transgenic fish. To increase the fluorescent intensity of the reporters, we crossed homozygote adult fish such that the embryo would carry 2 alleles of the reporters. The embryos were kept in 28°C for 4 hours, and then moved to 23°C. The embryos would typically arrive at 10 somite stage after 22-23 hours, at which we began the dissection. We dechorionated the embryos with 20mg/ml of pronase in embryo water. After the chorions were removed, we dissected the trunk region of about 200 embryos, using the same method described in the dual pipette aspiration assays. The dissection process typically took 1-2 hours. During this time, we kept the dissected tissues in 4°C. After the completion of dissection, dissected tissues were resuspended in 1ml FACSMax (Genlantis No. T200100) by manual pipetting to dissociate the cells and remove the yolk. Dissociated cells were spun down with pre-cooled centrifuge at 400G for 3 minutes and resuspended again in FACSMax. After the second treatment with FACSMax, cells were spun down and resuspended in pre-cooled PBS. Cells were sorted using a Sony SH800Z cell sorter, with a 100 μm chip. We performed the sort using the yield mode to increase the number of cells collected. At the end of the sorting session, sorted cells were examined by flow cytometry to verify purity. After the sorting was completed, sorted cells were spun down and the cell pellets were stored in trizol at −80°C. RNA was extracted automatically with QIAcube (Qiagen). We submitted 2 independent samples for each cell type (6 total) for RNA-seq. Each sample contains 2.4 ng of purified mRNA and was either obtained from one FACS sorting session or pooled from two different FACS sorting sessions. The RNA-seq library was prepared following the PrepX SPIA RNA-seq library protocol (IntegenX). Paired-End sequencing of the library was ran on a HiSeq 2500 system (Illumina).

## SUPPLEMENTARY DATA

**Figure S1.**
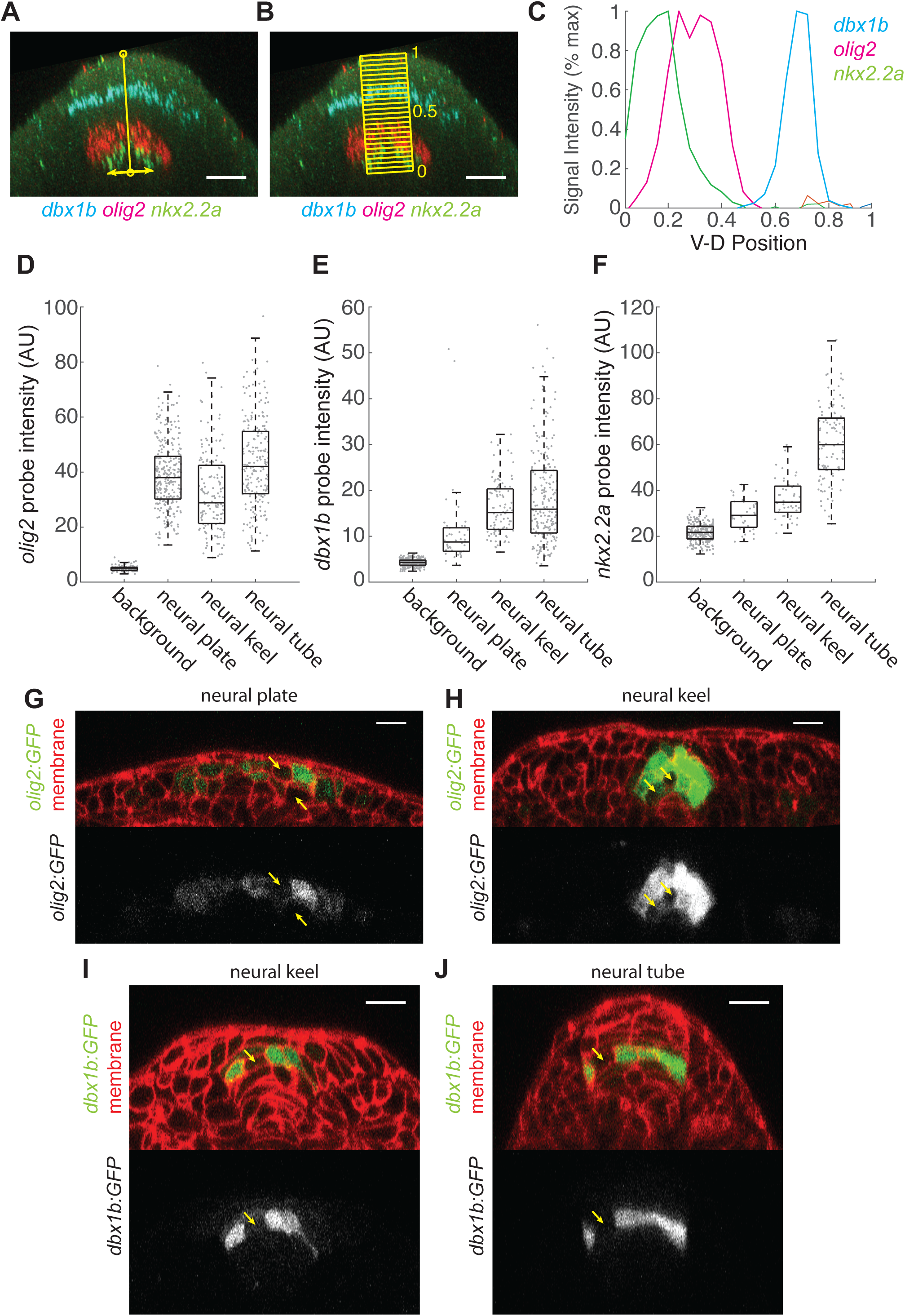
related to Figure 1. (A-C) Illustration of the analysis method to obtain the ventral-to-dorsal intensity profile of gene expression in the neural tube. For each cross-section, the ventral-most point of the neural tube and the dorsal-most point of the neural tube were manually selected, defining a ventral-to-dorsal axis (A). The width of the bounding box was estimated by the width of the p3 domain, typically spans 20 µm. 24 boxes were drawn along the ventral-to-dorsal axis with the pre-defined width (B). The average fluorescent intensity within each box in each fluorescent channel was calculated and plotted along the ventral-to-dorsal axis (C). Scale bars in (A,B) were 20 µm. (D-F) Estimation of single cell fluorescent intensity of *in situ* HCR probes in the (D) *olig2*, (E) *dbx1b*, (F) *nkx2.2a* positive domains. Each spot represents a 3.5 * 3.5 * 3.5 µm^3^ region in the fate marker positive domains. Background intensities were estimated by placing the volume in fate marker negative regions within the spinal cord tissues. The boxes within the box plot represent 25^th^, 50^th^, and 75^th^ percentile of the data. N denotes the number of embryos and n denotes the number of measurements in each condition. (D) *olig2* intensity, background (N=3, n=95), neural plate (N=3, n=277), neural keel (N=3, n=151), neural tube (N=3, n=212) (E) *dbx1b* intensity, background (N=3, n=277), neural plate (N=3, n=46), neural keel (N=3, n=95), neural tube (N=3, n=208), (F) *nkx2.2a* intensity, background (N=3, n=236), neural plate (N=4, n=32), neural keel (N=4, n=46), neural tube (N=3, n=104).). (G-J) Examples of fate-marker-negative cells mixed with the fate-marker-positive domains. (G,H) Yellow arrows point to individual *olig2:GFP* negative cells mixed within the *olig2:GFP* positive domains at the (G) neural plate stage, or (H) neural keel stage. Each cross-section has 2 mixed cells defined by our metric. (I,J) Yellow arrows point to individual *dbx1b*1:GFP negative cells mixed within the *dbx1b:GFP* positive domains at the (I) neural keel stage, or (J) neural tube stage. Each cross-section has 1 mixed cell defined by our metric. For (G-J), scale bars are 20 µm.

**Figure S2.**
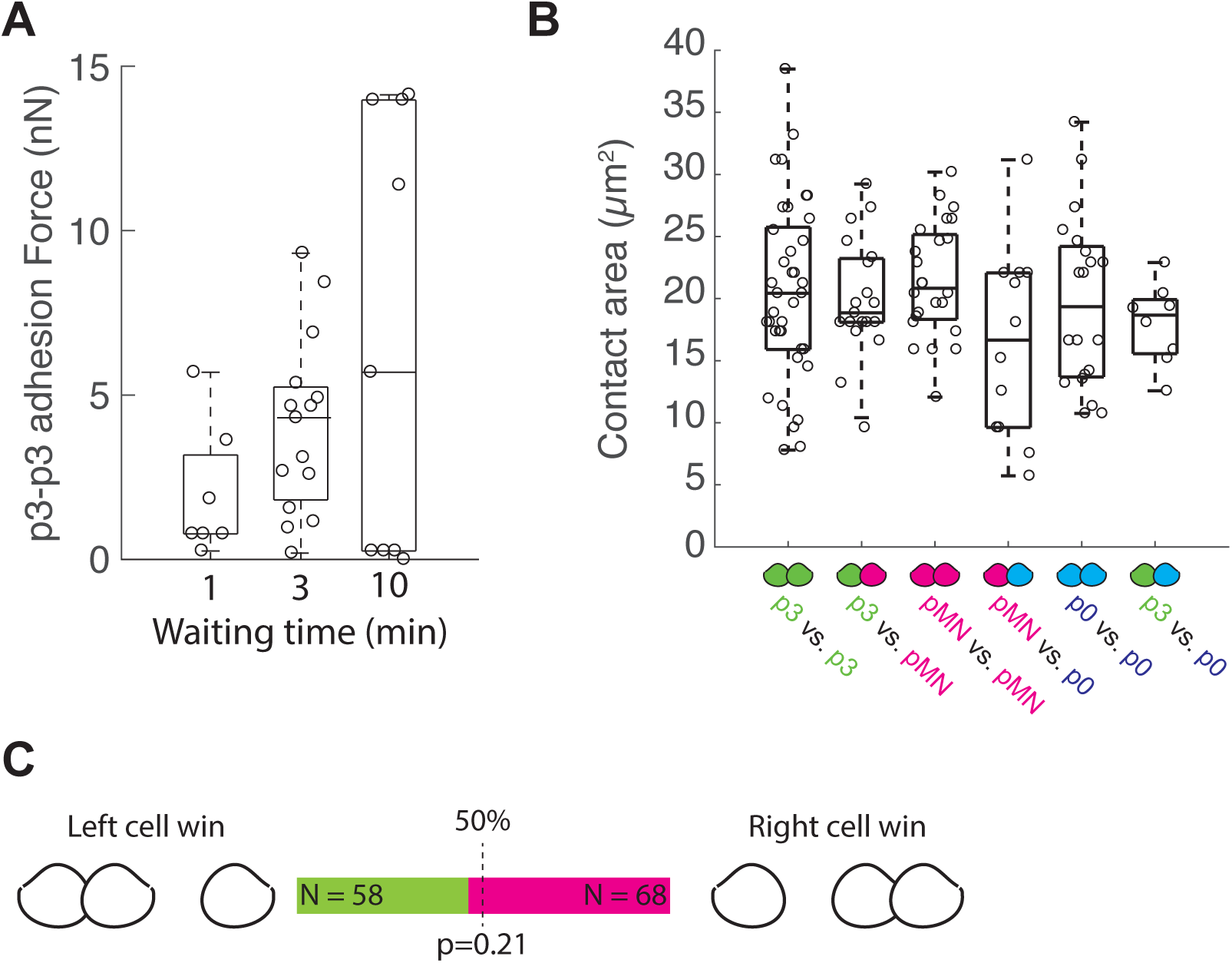
related to Figure 2. (A) Measurement of adhesion force between p3-p3 homotypic doublets under different duration of contact formation. Waiting time 1 minutes (N=3, n=9), 3 minutes (n=7, N=18), 10 minutes (N=3, n=9). The boxes within the box plot represent 25^th^, 50^th^, and 75^th^ percentile of the data. (B) Contact area of the six doublet types prior to pulling of micropipette. None of the doublet type has significant difference of contact area with other doublet types. p3-p3 (N=9, n= 37), p3-pMN (N=5, n=19), pMN-pMN (N=6, n=24), pMN-p0 (N= 3, n=12), p0-p0 (N=5, n=20), p3-p0 (N=2, n=8). The boxes within the box plot represent 25^th^, 50^th^, and 75^th^ percentile of the data. (C) No significant difference for the middle cell in the triplet to adhere to the left or the right cell of the triplet. P-value is determined by binomial test.

**Figure S3.**
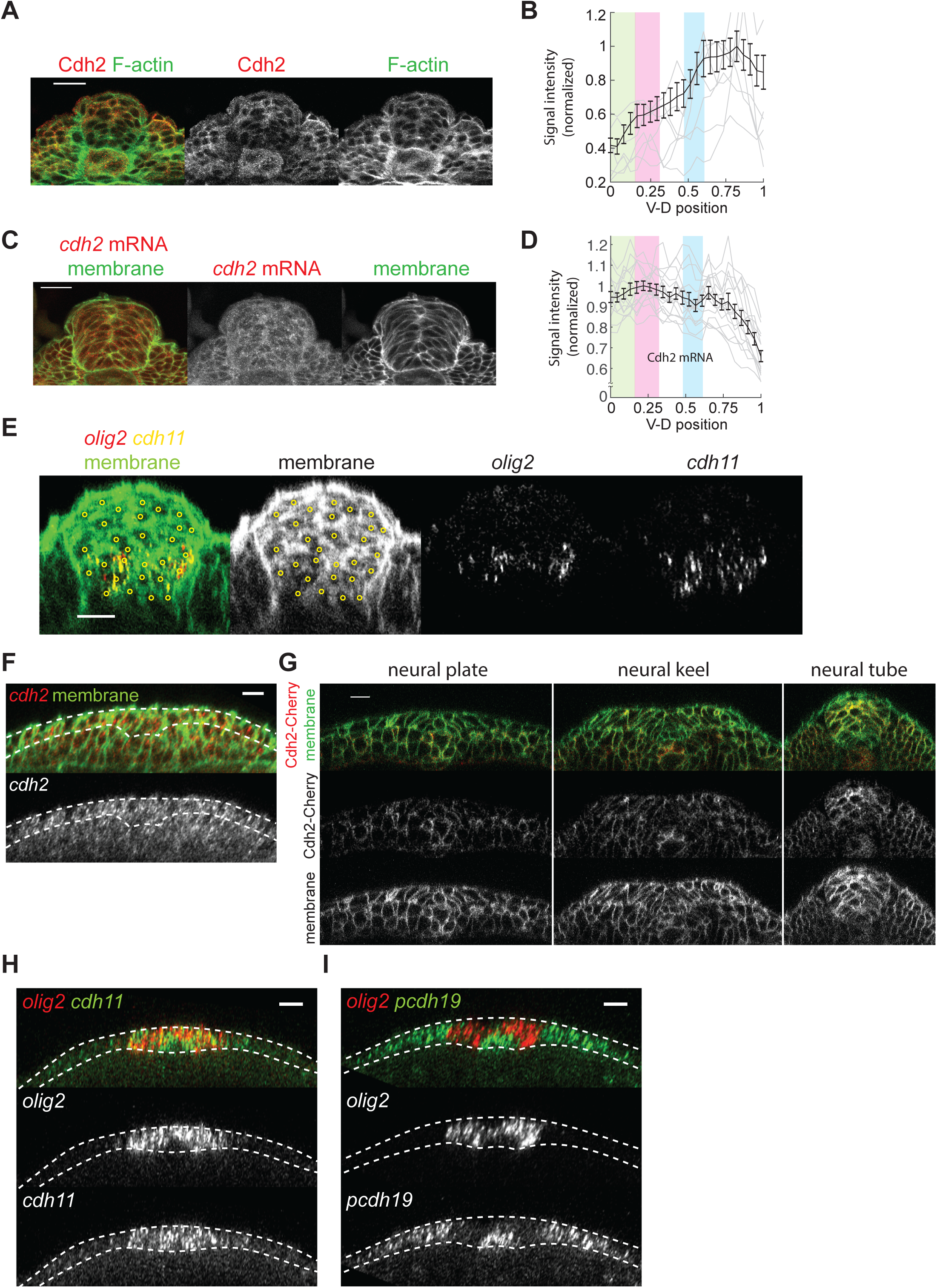
related to Figure 4. (A) Immunostaining of Cdh2 in sectioned neural tube of 10 somite stage embryos. F-actin was co-stained with phalloidin. (B) An average ventral-to-dorsal intensity profile of Cdh2 in 28 sectioned neural tube from 2 independent experiments. Gray traces in the background are selected single intensity profiles. Error bars are standard error of the mean. (C) *in situ* HCR of *cdh2* in sectioned neural tube of 10 somite stage embryos expressing ubiquitous membrane-Citrine marker. (D) An average ventral-to-dorsal intensity profile of 15 sectioned neural tube from 1 independent experiment. Gray traces in the background are selected single intensity profiles. Error bars are standard error of the mean. (E) A cross-section of a confocal z-stack in a transgenic embryo *Tg(actb2:memCitrine-Citrine)*, stained for *in situ* HCR probes against *olig2* and *cdh11*. Open yellow circles were manually defined as the center of cells for single cell analysis of gene co-expression. (F) A cross-section of a confocal z-stack of a *Tg(actb2:memCitrine-Citrine)* embryo at the neural plate stage. The embryo was stained with *in situ* HCR probes against *cdh2*. (G) Three cross-section snapshots of a 4-D time-lapsed confocal movie of a *Tg(actb2:memCitrine-Citrine)*; *TgBAC(cdh2:cdh2-Cherry)* embryo. The raw fluorescent intensity of the images is preserved for comparison of the Cdh2 level at the three stages. Initially, no clear differences of Cdh2 level can be observed across the neural plate. At the neural keel and neural tube stages, the Cdh2 level at the lateral/dorsal side increases significantly comparing to the neural plate stage, while the Cdh2 level at the medial/ventral side stays lower. Scale bar is 20 µm. (H) A cross-section of a confocal z-stack of an embryo at the neural plate stage. The embryo was stained with *in situ* probes against *olig2* (red) and *cdh11* (green). (I) A cross-section of a confocal z-stack of an embryo at the neural plate stage. The embryo was stained with *in situ* probes against *olig2* (red) and *pcdh19* (green).

**Figure S4.**
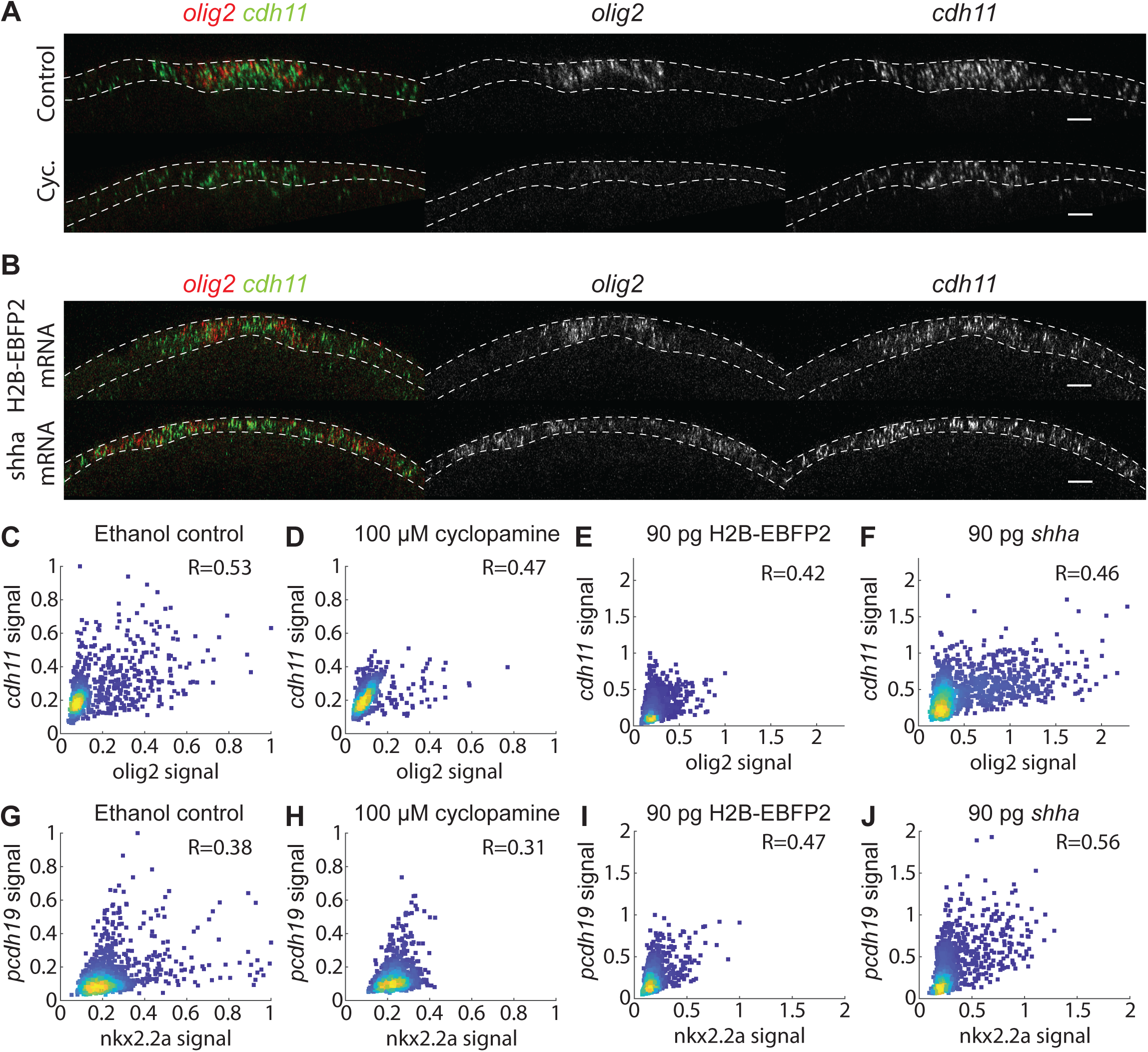
related to Figure 7. (A-B) Cross-sections of neural plate stage embryos treated with (A) ethanol vehicle control, or 100 µM cyclopamine, or (B) injected with H2B-EBFP2 or *shha* mRNA, and stained with *in situ* HCR probes against *olig2* and *cdh11*. Each cross-section is a maximal intensity projection of a 1.2 µm wide slab. Image contrast was adjusted for better visualization. Scale bars are 20 µm. (C-J) Single cell co-expression analysis of *olig2* versus *cdh11* (C-F) and *nkx2.2a* versus *pcdh19* (G-J) in embryos perturbed with Shh signaling. All axes were normalized to the maximum intensity of the control conditions. (C, D, G, H) Embryos were treated with ethanol vehicle control (C,G) or 100 µM cyclopamine (D,H), beginning at 4 hours post fertilization and fixed at 10 somite stage for multiplex *in situ* HCR staining. (E, F, I, J) Embryos were injected with 90 pg of mRNA of H2B-EBFP2 (E,I) or *shha* (F, J) at 1 cell stage and fixed at 10 somite stage for multiplex *in situ* HCR staining.

**Movie S1, related to Figure 1:**

Live imaging of a transgenic zebrafish embryo carrying fluorescent reporter *TgBAC(olig2:GFP); Tg(actb2:mem-Cherry)* to visualize the formation of the *olig2*-positive pMN domain. The tracked cell is labeled with yellow dot. Starting from the neural plate stage, we tracked a *olig2:GFP* positive cell found at 3 cell diameter (>40 µm) away from the rest of the *olig2:GFP* positive cells, mixing with the *olig2:GFP* negative cells. The tracked cell joins the remaining *olig2:GFP* positive cells during the transition from the neural plate to the neural keel stage. However, cell division takes place soon after the tracked cell joins the pMN domain, and one of the two divided cells is displaced away from the pMN domain. The tracked sister cell remains connected with the pMN domain while the other sister cell eventually rejoins the pMN domain. Finally, at the neural tube stage, the *olig2* positive cells form one cohesive stripe-like domain. This movie highlights the initial mixing pattern of the *olig2:GFP* cells and the detrimental effect of cell division for pattern cohesion. Scale bar of the movie is 20 µm and the image is taken every 6 minutes.

**Movie S2, related to Figure 1:**

Live imaging of a transgenic zebrafish embryo carrying fluorescent reporter *TgBAC(olig2:GFP); Tg(actb2:mem-Cherry)* to visualize the formation of the *olig2*-positive pMN domain. The tracked cell is labeled with yellow dot. The tracked cell was positioned at the tip of the wedge-shaped pMN domain during the neural keel stage, and it stayed connected with the pMN domain throughout this deformation process. Eventually, the tracked cell returned to a more ventral position and became part of a cohesive stripe of the pMN domain at the neural tube stage. Scale bar is 20 µm and the image is taken every 6 minutes.

**Movie S3, related to Figure 1:**

Live imaging of a transgenic zebrafish embryo carrying fluorescent reporter *TgBAC(dbx1b:GFP); Tg(actb2:mem-mCherry2)* to visualize the formation of the *dbx1b*-positive p0 domain. The one cell tracked throughout the movie is labeled with yellow dot. The *dbx1b:GFP* signal is visible shortly after the neural plate stage at the beginning of the movie. At the neural keel stage, the *dbx1b* positive cells are found on either side of the neural keel, with *dbx1b* negative cells in between. The *dbx1b* positive cells continue to migrate dorsal-medially and make contacts with the *dbx1b* positive cells from the other side of the neural keel. The tracked *dbx1b* positive cell divided as it approaches the midline at the neural keel stage, and the tracked sister cell crossed the midline to reach the right side of the neural tube. Eventually the *dbx1b:GFP* positive cells settle in a stripe-like p0 domain. Scale bar is 20 µm and the image is taken every 6 minutes.

**Movie S4, related to Figure 1:**

Live imaging of transgenic zebrafish embryo carrying fluorescent reporter *TgBAC(nkx2.2a:mem-GFP); Tg(actb2:mem-mCherry2)* to visualize the formation of the *nkx2.2a*-positive p3 domain. The contrast of the movie is adjusted to highlight the GFP signal and filtered by a median filter to remove noise. The one cell tracked throughout the movie is labeled with yellow dot. The *nkx2.2a:mem-GFP* signal became visible at the neural keel stage in the tracked cell in this movie. Initially, we only see one *nkx2.2a* positive cell in the whole cross section, as the *nkx2.2a* positive cells were scattered along the anterior-posterior axis. The tracked cell then divided into two sister cells that eventually settled into either side of the medial floor plate. Scale bar is 20 µm and the image is taken every 7 minutes.

**Movie S5, related to Figure 2:**

Movie of a dual pipette aspiration assay to separate 2 p3 cells, as identified by the fluorescent reporter *TgBAC(nkx2.2a:mem-GFP)*. The p3-p3 doublet was allowed to adhere for more than 3 minutes before the pulling starts. This movie shows a pulling sequence of 40→60→80→100 Pa. Since the doublet breaks at 100 Pa, we recorded the separation pressure of 90 Pa from the average of 80 and 100 Pa.

**Movie S6, related to Figure 2:**

Movie of a triplet assay showing homotypic preference of p0 cells (green only) versus pMN cell (green and magenta). The p0 cell is obtained from dissociation of transgenic embryos carrying *TgBAC(dbx1b:GFP).* The pMN cell is obtained from dissociation of transgenic embryos carrying *TgBAC(olig2:GFP).* The *TgBAC(olig2:GFP)* embryos were injected with 1ng of Texas Red conjugated 3kD dextran in this experiment. The initial assembly of the triplet was shown in the first part of the movie. The left cell is a *olig2:GFP* positive pMN cell, while the middle and right cells are *dbx1b:GFP* positive p0 cells. The triplet was allowed to form adhesion for 3.5 minutes. In the second part of the movie, the pulling of the triplet was shown. The right pipette was pulled towards the right of the stage with a controlled motor at a speed of 20 µm/sec. In this example, the p0-p0 homotypic contact remained while the p0-pMN contact broke during the pulling, suggesting the homotypic preference of the p0 cells against the pMN cell.

**Movie S7, related to Figure 6:**

Live imaging of a *cdh11*-morphant embryo carrying fluorescent reporter *TgBAC(olig2: GFP); Tg(actb2:mem-mCherry2)* to visualize the formation of the *olig2*-positive pMN domain. The one cell tracked throughout the movie is labeled with yellow dot. The tracked cell was positioned at the tip of the wedge-shaped pMN domain, and detached from the rest of the pMN domain during the neural keel stage. Eventually, the *olig2:GFP* positive cell mislocalized to the dorsal side of the pMN domain at the neural tube stage. This is different from the behavior of the *olig2:GFP* cell in the wildtype embryo, as shown in Movie S2.

