## Supplementary figures and images for "An adhesion code ensures robust pattern formation during tissue morphogenesis"

### FigureS1

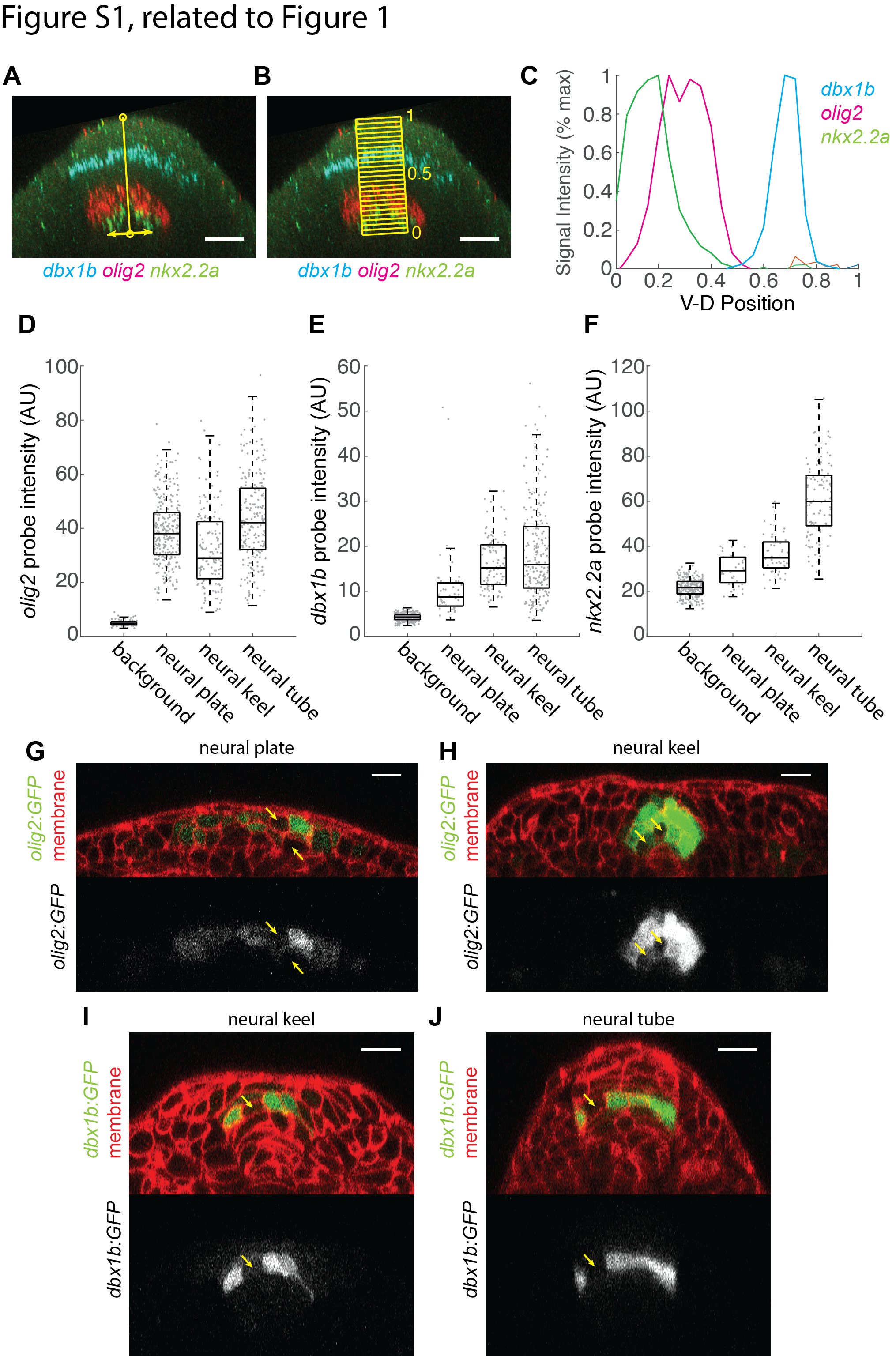

### FigureS2

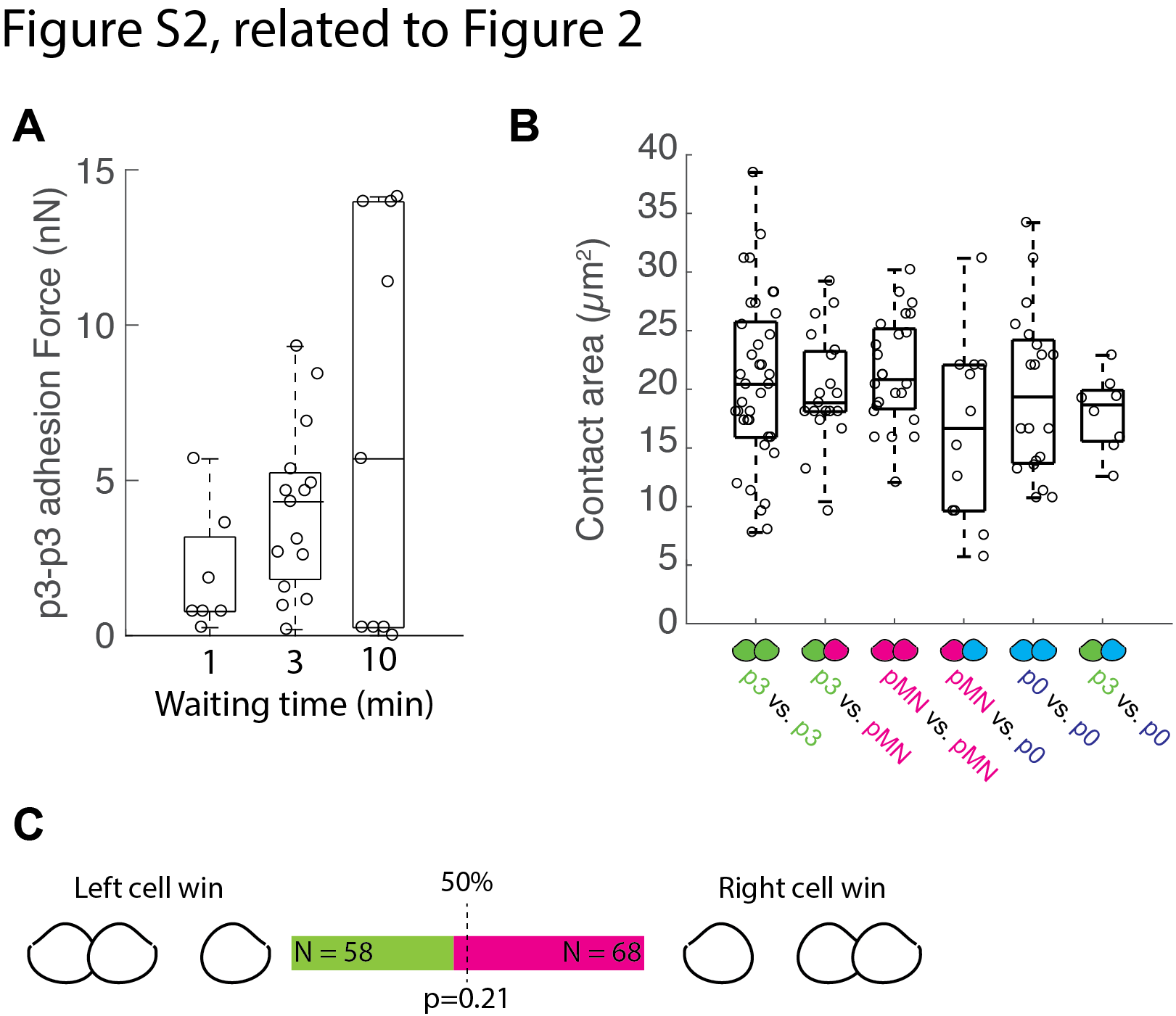

### FigureS3

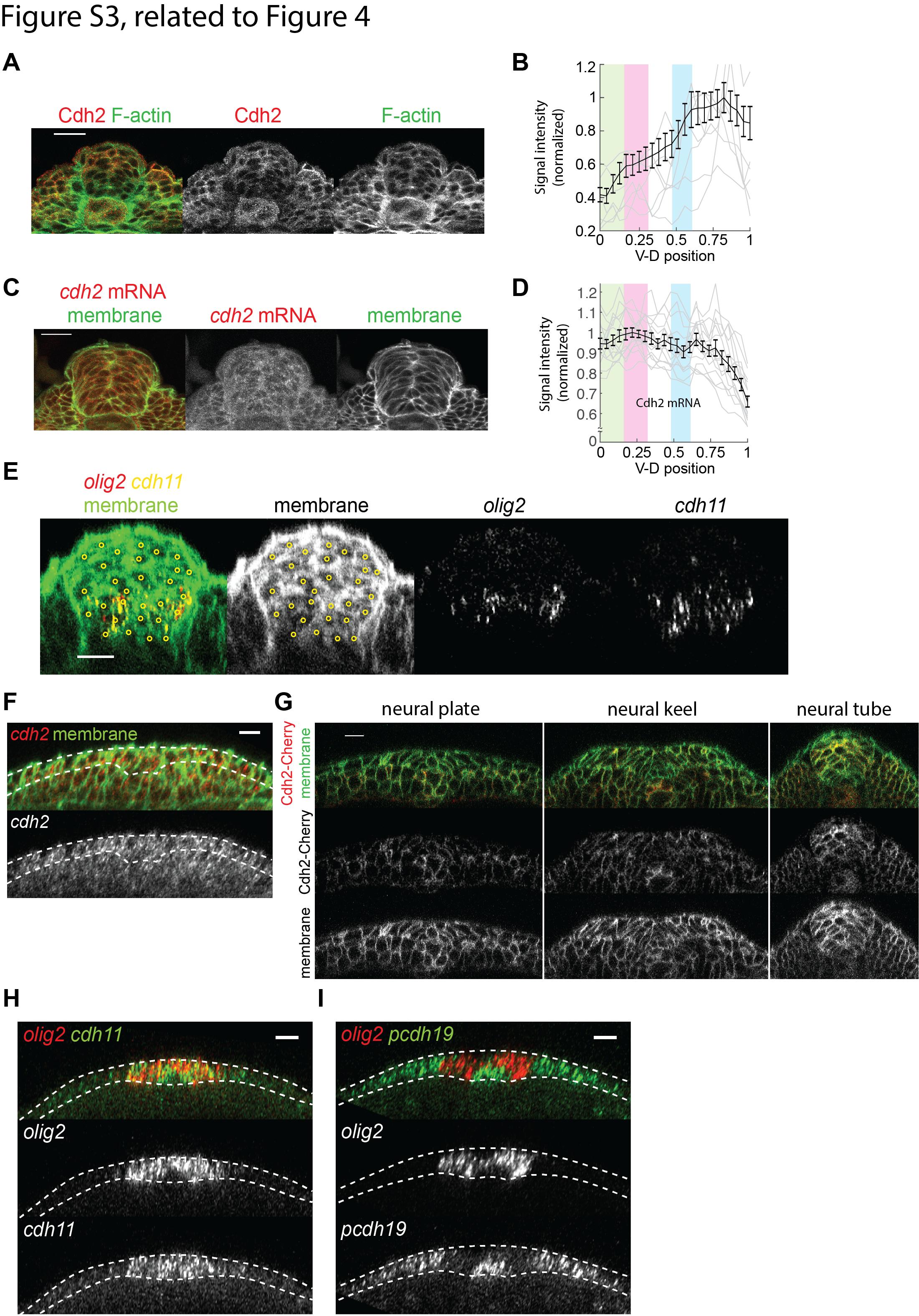

### FigureS4

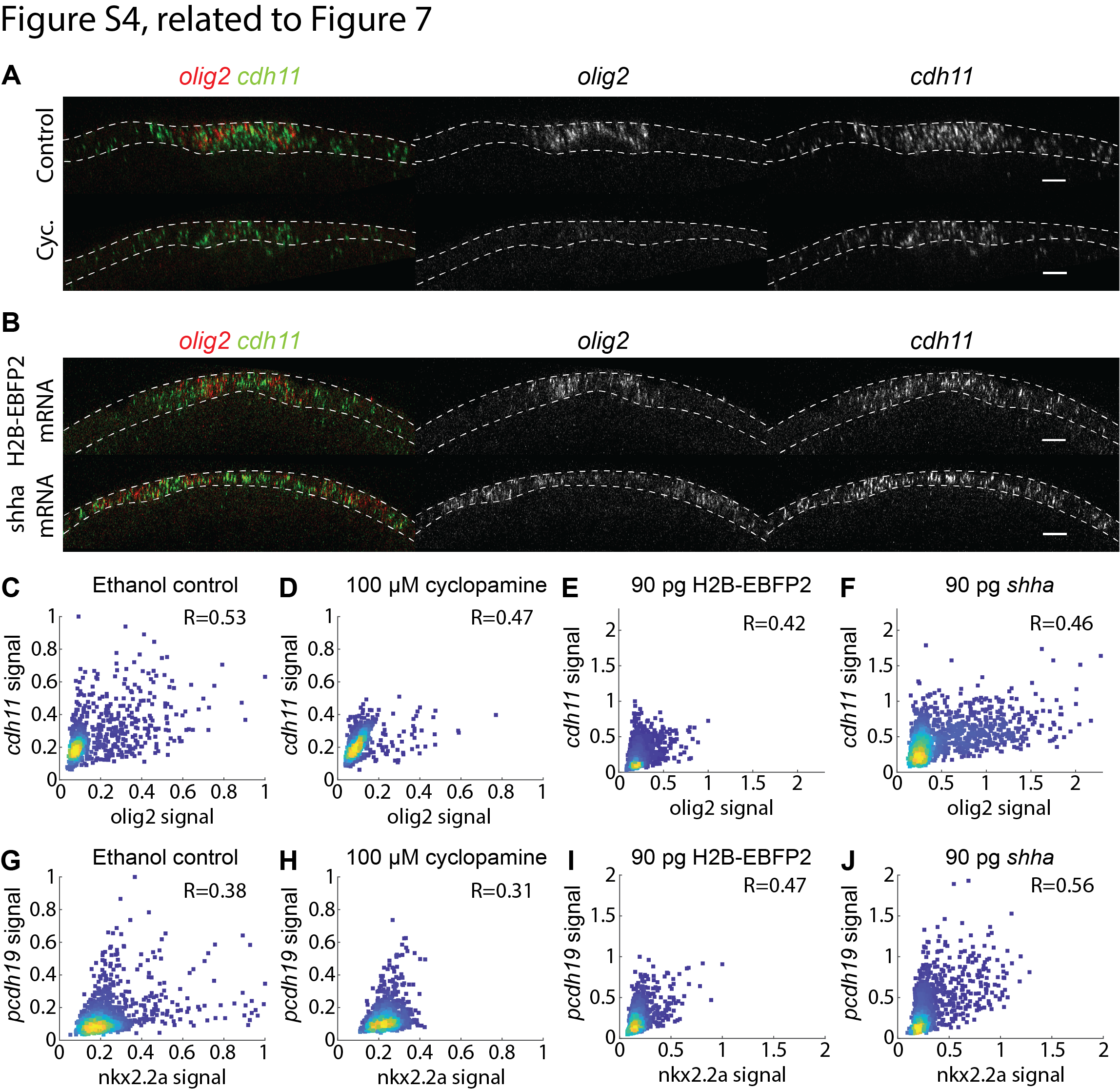
